# Epilepsy kinase CDKL5 is a DNA damage sensor which controls transcriptional activity at DNA breaks

**DOI:** 10.1101/2020.12.10.419747

**Authors:** Taran Khanam, Ivan Muñoz, Florian Weiland, Thomas Carroll, Barbara N Borsos, Vasiliki Pantazi, Meghan Slean, Miroslav Novak, Rachel Toth, Paul Appleton, Tibor Pankotai, Houjiang Zhou, John Rouse

## Abstract

Mutation of the *CDKL5* kinase gene leads to the seizure-prone neurodevelopmental condition CDD (CDKL5 deficiency disorder) and is the most common genetic cause of childhood epilepsy. However, the phospho-targets and roles of CDKL5 are poorly understood, especially in the nucleus. We reveal CDKL5 as a sensor of DNA damage in actively transcribed regions of the nucleus, which phosphorylates transcriptional regulators such as Elongin A (ELOA) on a specific consensus motif. Recruitment of CDKL5 and ELOA to DNA damage sites, and subsequent ELOA phosphorylation, requires both active transcription and synthesis of poly–ADP ribose to which CDKL5 can bind. Critically, CDKL5 is essential for transcriptional control at DNA breaks. Therefore, CDKL5 is a DNA damage-sensing regulator of transcription, with implications for CDKL5-related human diseases.

**One sentence summary:** CDKL5 is a DNA damage-sensing kinase that modulates transcriptional activity near DNA breaks.

---

The protein kinases ATM, DNA-PK and ATR sense and transduce DNA damage signals, triggering a pleiotropic series of protective reactions collectively known as the DNA damage response (DDR) which prevents genome instability and disease (*1*). With the aim of expanding the repertoire of DDR kinases, we set out to find other kinases that can sense DNA damage. U–2–OS cells stably expressing mCherry–FAN1 which marks DNA damage sites, were transfected with GFP–tagged kinases individually, starting with the CMGC branch of the human kinome (Fig. S1A). BrdU–sensitized cells were laser micro-irradiated to induce a pleiotropic range of DNA lesions along a track in the nucleus. This approach revealed CDKL5 as a DNA damage sensing kinase (Fig. 1A, S1A). *CDKL5* mutations are one of the most common genetic causes of epilepsy in children (*2*), and they can lead to the severe, seizure prone neurodevelopmental disorder CDD (*3*), as well as milder syndromes (*4*). In the cytosol, CDKL5 phosphorylates microtubule regulators (*5, 6*), but the nuclear roles and targets of CDKL5 remain elusive. This prompted us to investigate how CDKL5 recognizes DNA damage, to find its nuclear targets and explore roles in DDR.

**Figure 1.**
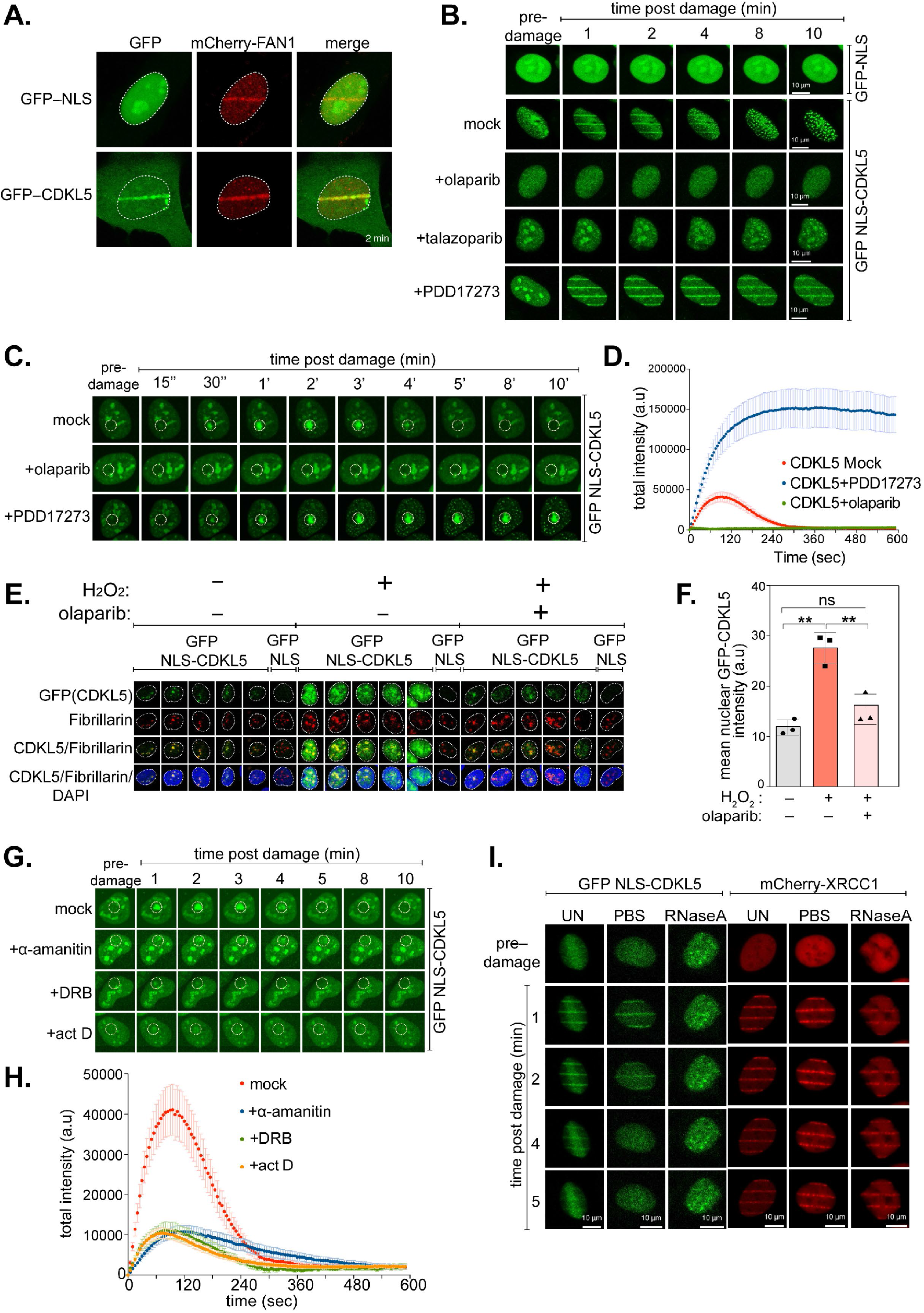
CDKL5 senses DNA damage in actively transcribed regions. **A.** BrdU–sensitized U–2–OS Flp–In T–REx stably expressing mCherry-FAN1 and GFP–NLS or GFP–CDKL5 (no NLS) were line micro-irradiated and imaged after 2min. **B.** Same as A. except that cells stably expressing GFP–NLS–CDKL5 were pre–incubated with DMSO (mock), olaparib (5 μM), talazoparib (50 nM) or PDD00017273 (0.3 μM) for 1 h prior to micro–irradiation. One of three independent experiments is shown. **C.** Same as B. except that cells were spot micro–irradiated (405 nm). **D.** Quantitation of spot intensity in C. Data represent the mean ± SEM of two independent experiments; > 50 micro–irradiated cells per point. **E.** Cells subjected to the workflow in Fig. S1C were detergent–extracted and fixed before staining with anti–GFP or fibrillarin (nucleoli). **F.** Quantification of the detergent–insoluble GFP–NLS–CDKL5 signal (minus nucleolar signal). The mean ± SD from three biological experiments is shown. Statistical significance was assessed by one-way-ANOVA-test. Asterisks ** indicate *P*–value of <0.01; ns – not significant. **G, H.** Effect of transcription inhibitors on CDKL5 recruitment after spot micro-irradiation. **I.** Stable cell lines were permeabilized and incubated with RNase A or PBS before irradiation and imaging.

CDKL5 recruitment to sites of micro–irradiation was rapid and transient (Figs. 1A–D, Movies S1, S2), reminiscent of proteins that bind poly–ADP ribose (PAR) generated by DNA damage–activated poly–ADP ribose polymerases (PARPs) (*7*). Accordingly, CDKL5 recruitment was blocked by PARP inhibitors olaparib and talazoparib (Figs. 1B–D) or by *PARP1* disruption (Fig. S1B), but prolonged by PDD00017273, an inhibitor of PARG (poly–ADP ribose glycohydrolase) which delays PAR degradation (Fig. 1B–D; Movies S1, S2) (*8*). CDKL5 was also retained on damaged chromatin after exposure to H_2_O_2_, an inducer of DNA breaks (Figs. 1E, F; S1C). Retention was prevented by olaparib, whereas nucleolar retention seen in undamaged cells was unaffected (Figs. 1E, F). Together, these data indicate that CDKL5 recruitment to DNA breaks requires synthesis of PAR, presumably by direct PAR binding although no PAR-binding motifs were detected bioinformatically. However, a series of N-terminal and C-terminal deletion constructs revealed a region between amino acids 530 and 730 necessary for CDKL5 recruitment (Figs. S2A, B), and the region 530–680 of CDKL5 was sufficient for recruitment (Figs. S2C, D). As shown in Figs. S2D–F, recombinant CDKL5 fragments corresponding to this region bound to PAR *in vitro* (Fig. S2E, F), strongly suggesting that CDKL5 senses DNA breaks by binding PAR. Accordingly, PAR was detected in CDKL5 precipitates, and vice versa, after exposure of cells to H_2_O_2_ (Figs. S2G,H).

These data show that PAR formation is required for CDKL5 recruitment, but we discovered unexpectedly that it is not sufficient. We found that the transcription inhibitors actinomycin D or α–amanitin which inhibit RNA polymerases (RNAPs) I and II, or DRB which blocks elongating RNAP II, abrogated the recruitment of CDKL5 to micro-irradiation sites. PAR synthesis and recruitment of the PAR-binding protein XRCC1 or FAN1 was unaffected (Figs 1G,H; S3A–D). Moreover, incubation of permeabilized cells with RNaseA abolished micro-irradiation tracks formed by CDKL5, but not XRCC1 or FAN1 (Fig. 1I). Therefore, CDKL5 is recruited to DNA breaks at sites of active transcription.

The data above suggested that nuclear targets of CDKL5 may be involved in transcriptional control. To identify nuclear CDKL5 targets specifically, we expressed CDKL5 wild–type (WT) or a K^42^R kinase–dead (KD) mutant (*5*) exclusively in the nucleus of *CKDL5-disrupted* U–2–OS cells by adding an artificial nuclear localization signal (NLS) (Fig. S4A–C). We next compared the phosphoproteome of the two cell populations, after exposure to H_2_O_2_ to induce CDKL5 retention at DNA breaks (Fig. 2A, S4D, Table S1). As well as CDKL5 itself and MAP1S and EB2 (known cytosolic substrates), screening revealed a range of proteins bearing phospho–sites that were higher in abundance in CDKL5^NLS^–WT cells compared with CDKL5^NLS^–KD cells (>1.5–fold, p<0.05) (Figs. 2B, C; Table S1). Strikingly, the phospho–acceptor [S/T] residue in almost all of the putative nuclear CDKL5 substrates lies in sequence R–P–X–[**S/T**]– [A/G/P/S] (Fig. 2C), which represents a prerequisite consensus motif for CDKL5 target phosphorylation, in agreement with previous work (*5, 6*). Gene ontology (GO) analysis showed a striking enrichment of transcription regulators (Figs. 2D, E) including EP400, a chromatin-remodeling transcriptional activator (*9*), and TTDN1, mutated in a form of tricothiodystrophy (TTD), typically caused by failure in transcription–coupled DNA repair (*10*). Elongin A (ELOA), a transcriptional elongation factor and component of an E3 ligase complex which ubiquitylates RNAPII, was a top hit (*11*).

**Figure 2.**
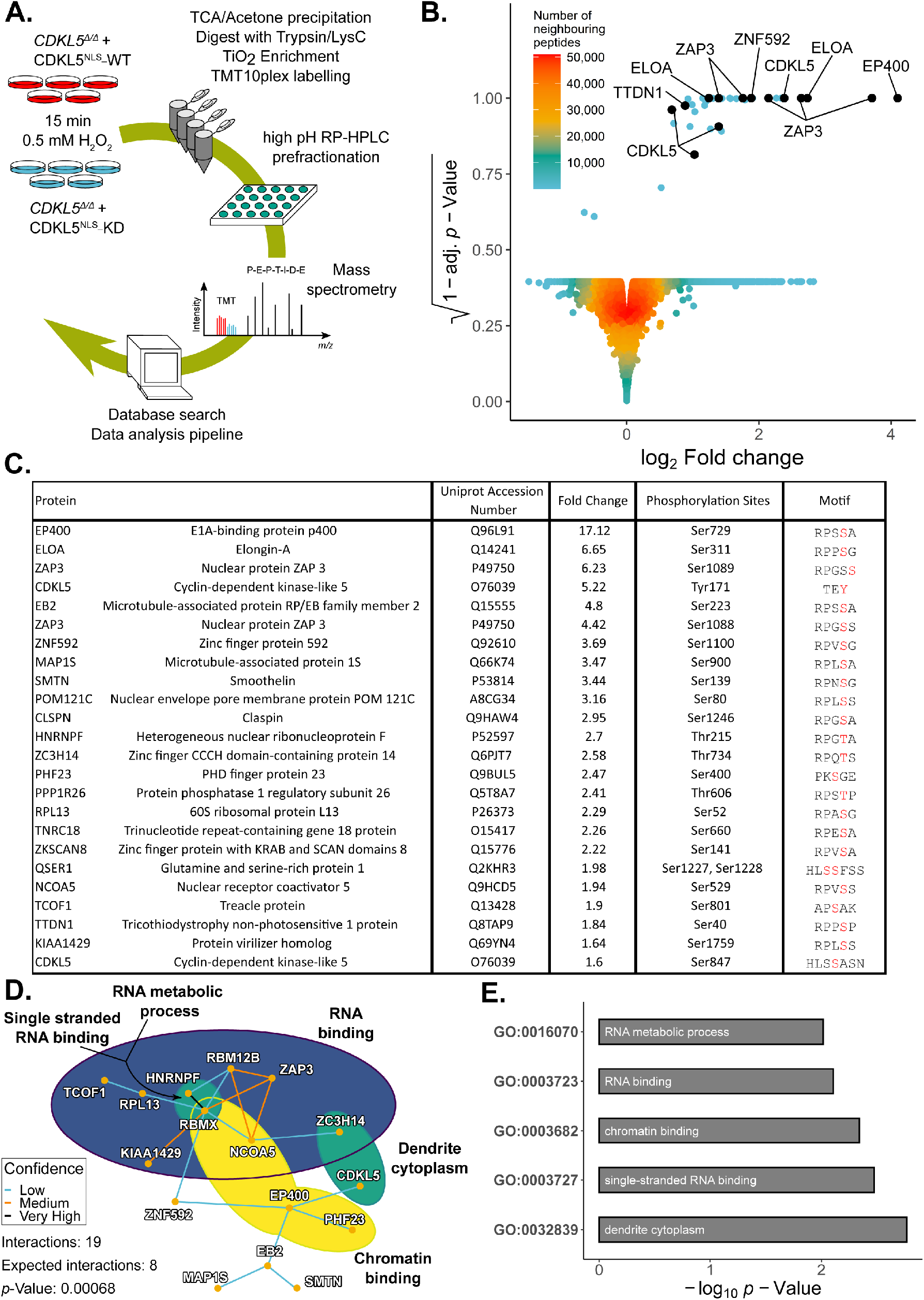
CDKL5 phosphorylates transcriptional regulators including ELOA. **A.** Quantitative phosphoproteomics workflow. **B.** “Sprinkler” plot of the mass spectrometry data from the experiment in A (see Table S1). Phosphorylated residues are highlighted in red. **C.** List of proteins containing phosphorylation sites more abundant in cells expressing CDKL5^NLS^–WT versus KD (fold change > 1.5; p-site probabilities > 0.6). **D.** Protein–protein interaction network of putative CDKL5 substrates from Table S1. Confidence levels are based on the STRING database v11.0 combined score with following bins: 150–400: low confidence (blue), 400–700: medium confidence (gold), 700–900: high confidence (not encountered in this dataset), >900: very high confidence (black). P–value was calculated as 0.00068. **E.** Analysis of GO terms. Significance cut–off was set as α = 0.01 with at least 2 proteins identified in the respective group.

Extracted ion chromatogram (XIC) analysis of phospho–peptides isolated from FLAG-tagged EP400 (pSer^729^), ELOA (pSer^311^) and TTDN1 (pSer^40^) confirmed CDKL5-dependent phosphorylation of these proteins in cells (Fig. S5A-C). Furthermore, CDKL5 robustly phosphorylated synthetic peptides corresponding to EP400 Ser^729^ and ELOA Ser^311^ demonstrating direct phosphorylation (Fig. S5D). To further investigate ELOA phosphorylation, we generated antibodies specific for phospho-Ser^311^. Co-expression with WT, but not KD, CDKL5 markedly increased Ser^311^ phosphorylation of FLAG-ELOA, but not an ELOA Ser^311^Ala mutant (Fig. 3A). Strikingly, CDD–associated CDKL5 mutations, which are located predominantly in the kinase catalytic domain (*12*), severely reduced CDKL5 activity towards ELOA pSer^311^, whereas a series of benign variants did not (Fig. 3B).

**Fig. 3.**
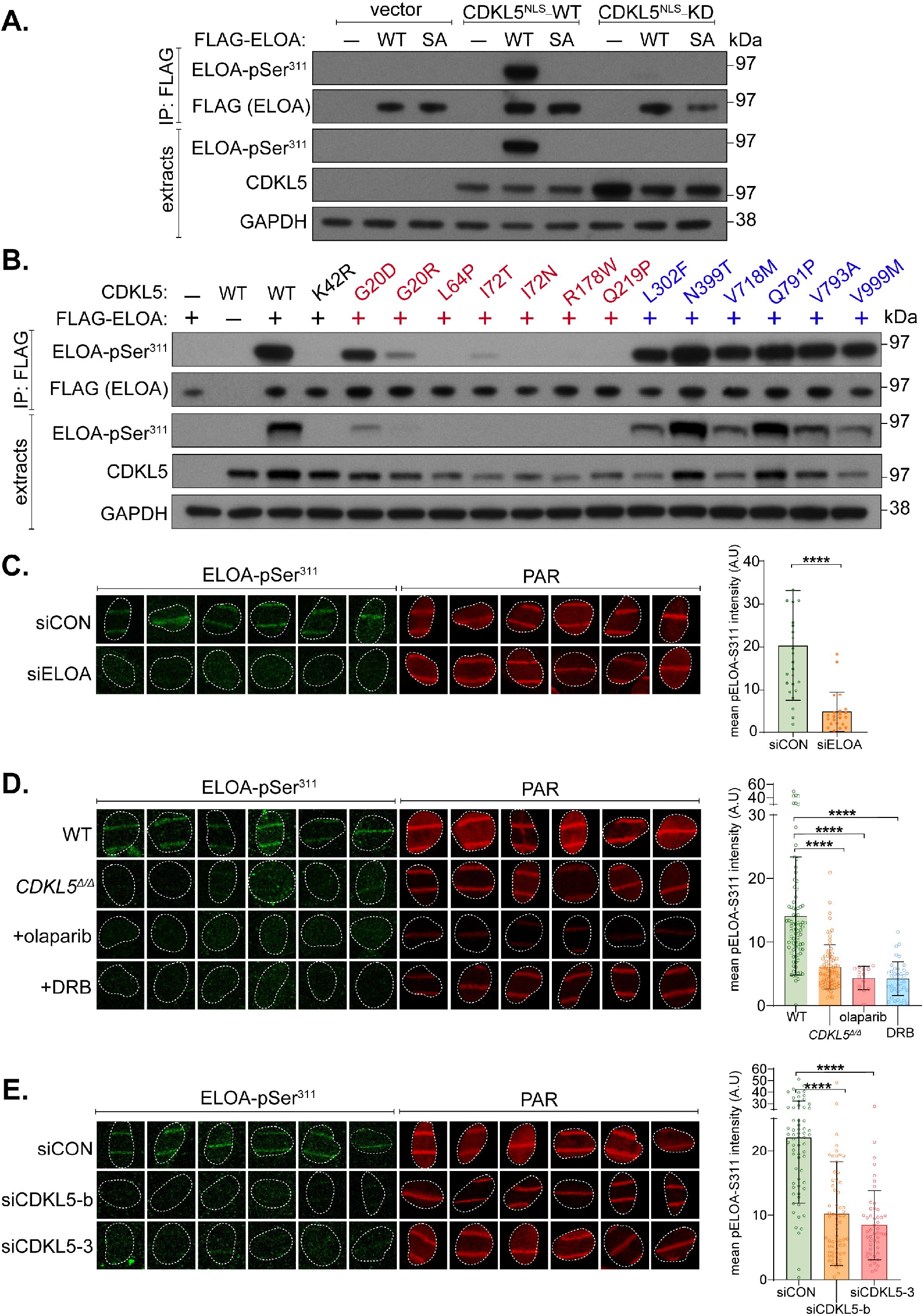
Phosphorylation of ELOA Ser^311^ on damaged chromatin by CDKL5. **A.** HEK293 cells were co-transfected with CDKL5 (wild type “WT” or kinase-dead “KD” K^42^R mutant) fused to an NLS, and FLAG-ELOA (wild type “WT” or a S^311^A mutant “SA”). Anti-FLAG precipitates or cell extracts were probed with the antibodies indicated. One of three independent experiments is shown. **B.** Same as A. showing a range of pathogenic (red) and benign (blue) CDKL5 variants. **C–E**. Wild type (WT), CDKL5–disrupted (CDKL5^*Δ/Δ*^) or siRNA–transfected cells were subjected to indirect immunofluorescence analysis with the indicated antibodies at laser tracks. Quantification of ELOA-pSer^311^ signal at the laser tracks is shown. Data represent mean ± SD. Statistical significance was assessed by one-way-ANOVA-test. Asterisks **** indicate *P*–values of <0.0001.

CDKL5-dependent phosphorylation of endogenous ELOA Ser^311^ was evident at sites of DNA damage. Signal intensity was reduced by depletion of ELOA (Fig. 3C) or by incubation of cells with lambda-phosphatase or the ELOA Ser^311^ phosphopeptide antigen (Fig. S6A), thereby confirming antibody specificity. ELOA phosphorylation was reduced by disruption or depletion of *CDKL5* (Figs. 3D, E), or by olaparib or DRB which block CDKL5 recruitment (Fig. 3D). We wondered if ELOA is recruited to DNA damage sites by a similar mechanism to the ELOA kinase. In agreement with this idea ELOA recruitment to micro-irradiation tracks was rapid, transient, and inhibited by olaparib, α-amanitin and DRB (Fig. S6B–D). Similar results were obtained for other nuclear CDKL5 substrates such as ZNF592 and ZAP3 (Fig. S6B-D) but not EP400 (data not shown). These data reveal CDKL5-dependent phosphorylation of substrates such as ELOA at DNA damage sites, involving a common mechanism of recruitment of both kinase and substrate.

The ontological enrichment for transcription regulators among the nuclear CDKL5 substrates suggested a role in transcriptional control at DNA damage sites. Breaks in genomic DNA silence adjacent genes (*13–15*), and we tested a role for CDKL5, first using a reporter system in which a cluster of FokI nuclease–mediated DSB, induced upstream of a doxycycline inducible reporter gene, silences the reporter cassette (Fig. S7A) (*13*). Depletion of CDKL5 weakens silencing of the reporter cassette, similar to depletion of ATM or ZMYND8 (Fig. 4A; Figs. S7A–C) (*13, 14*). Second, we took advantage of a system where inducible overexpression of the site-specific meganuclease I-PpoI that cuts 14–30 times in the human genome, results in silencing of genes that sustain DSBs within or nearby. As shown in Fig. 4B, CDKL5 depletion largely abolished the I–PpoI–induced silencing of *SLCO5a1* and *RYR2* genes reported previously (*15*).

**Fig. 4.**
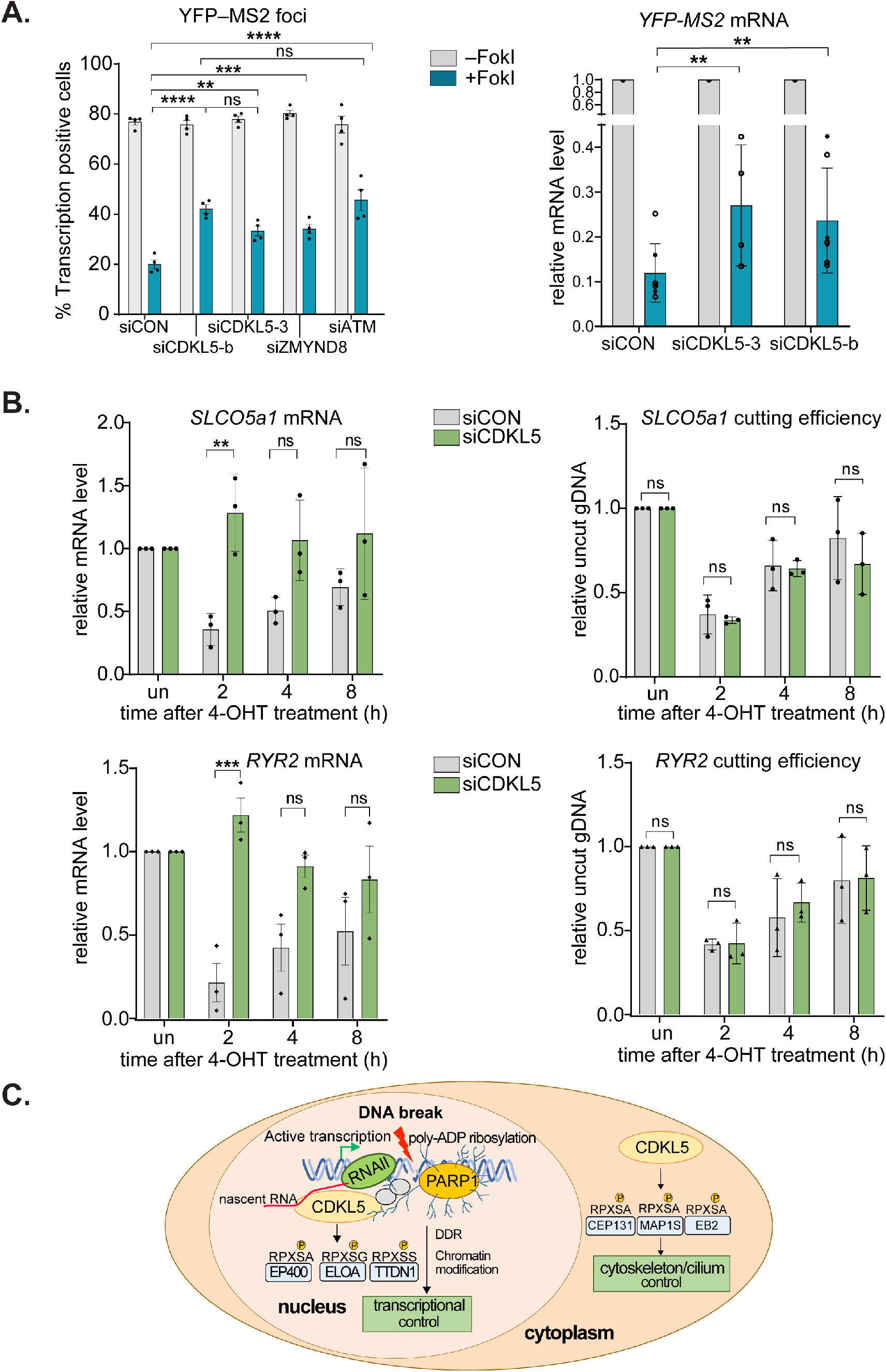
CDKL5 facilitates transcriptional silencing at DNA breaks. **A.** U–2–OS cells (263 IFII; Fig. S7A) were transfected with the siRNAs indicated. After addition of doxycycline, transcription was monitored in cells ± induction of FokI by quantification of YFP(−MS2) foci (*left*). 150 cells were analysed per condition per experiment. The mean ± SD from four independent experiments is shown. (*Right*) Quantitative RT–PCR analysis of YFP-MS2 mRNA. Data represent mean ± SD. **B**. Quantitative PCR with reverse transcription (qRT-PCR) analysis of *SLCO5a1* and *RYR2* expression levels (left) and cutting efficiencies (right) in U–2–OS HA-ER-I–PpoI cells depleted of CDKL5, at the times indicated after inducing I-PpoI. The mean ± SD from qPCR replicates of three independent experiments is shown Statistical significance for all the data was assessed by two-way-ANOVA-test **p<0.01, ***p<0.001 and ****p<0.0001; ns – not significant. **C.** Schematic diagram: CDKL5 functions in nucleus and cytosol.

Our study reveals that CDKL5 senses DNA breaks at sites of ongoing transcription, through binding to PAR, and switches off transcription nearby (Fig. 4C). How transcriptional activity at DSB is sensed by CDKL5 remains to be investigated. Once bound to damaged DNA, CDKL5 phosphorylates transcriptional regulators including ELOA, and facilitates the silencing of genes harboring DNA breaks; it is likely that phosphorylation of multiple CDKL5 substrates participates in silencing. For example, CDKL5 phosphorylation of ELOA could influence transcriptional elongation rates, but this remains to be ascertained. CDKL5 was recently linked to transcriptional control in a different context when it was reported to promote renal injury in mice exposed to toxic insults through up-regulating SOX9-dependent genes (*16*). We are interested in the possibility that CDKL5 controls transcription even in the absence of toxic insult, perhaps by sensing the transient DSBs induced by topoisomerases which are known to regulate transcription and impact on brain function (*17*). This will be interesting to investigate, especially as it could be particularly relevant to the pathogenesis of epilepsy and CDD.

## Acknowledgements

We thank the technical support of the MRC-PPU including the DNA Sequencing Service, Tissue Culture team, Reagents and Services team, and the PPU Mass-Spectrometry team. We also thank Fiona Brown and James Hastie for ELOA antibody production and purification. We’re grateful to Jessica Downs and Roger Greenberg for the U–2–OS cells harbouring the FokI silencing reporter, to Evi Soutoglou for U–2–OS HA-ER-I-PpoI cells, and to Nick Lakin for *PARP*-disrupted U–2–OS cells. We’re grateful to Graeme Ball for help with microscopy data analysis. We thank Luis Sanchez-Pulido and Chris Ponting for help with bioinformatic analyses. We are grateful to members of the Rouse lab for useful discussions. This work was supported by the Medical Research Council (grant number MC_UU_12016/1; TK, IM, JR) and the pharmaceutical companies supporting the Division of Signal Transduction Therapy Unit (Boehringer-Ingelheim, GlaxoSmithKline, and Merck KGaA).

---

**Fig. S1.**
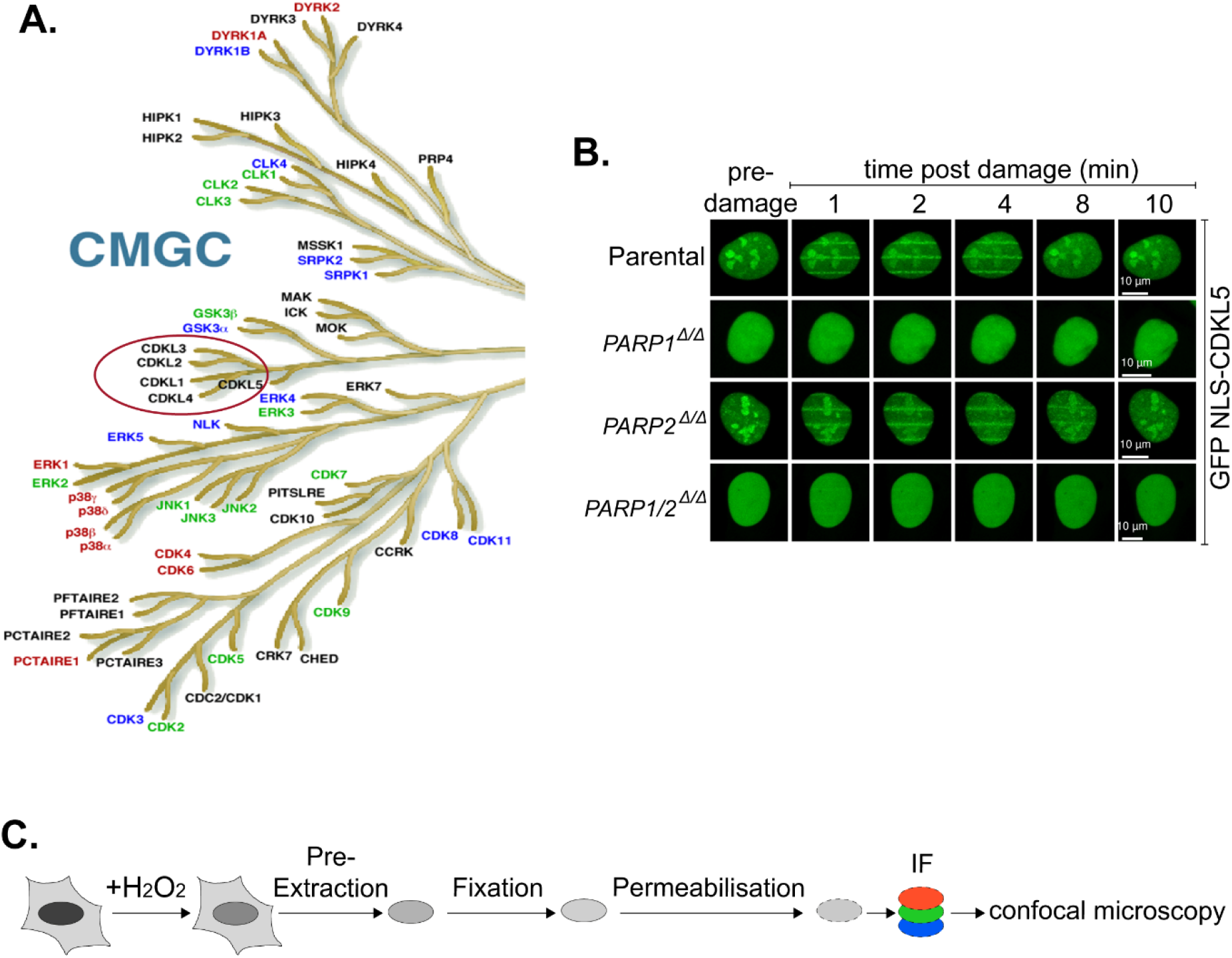
Recruitment of CDKL5 to sites of DNA damage. (**A**) Schematic diagram of the CMGC branch of the human kinome taken from the dendrogram made by Manning and colleagues (*1*). The CDKL family of kinases which includes CDKL5 is encircled in red. (**B**) BrdU–sensitized *PARP1*^Δ/Δ^, *PARP2*^Δ/Δ^, *PARP1/2*^ΔΔ^ cells, or parental U–2–OS cells transiently expressing GFP NLS–CDKL5 were subjected to 355 nm line micro–irradiation followed by time lapse imaging. Two independent experiments were performed, and one representative experiment is shown. (**C**) Diagram of the workflow for the chromatin retention experiments shown in Fig 1E, F.

**Fig. S2.**
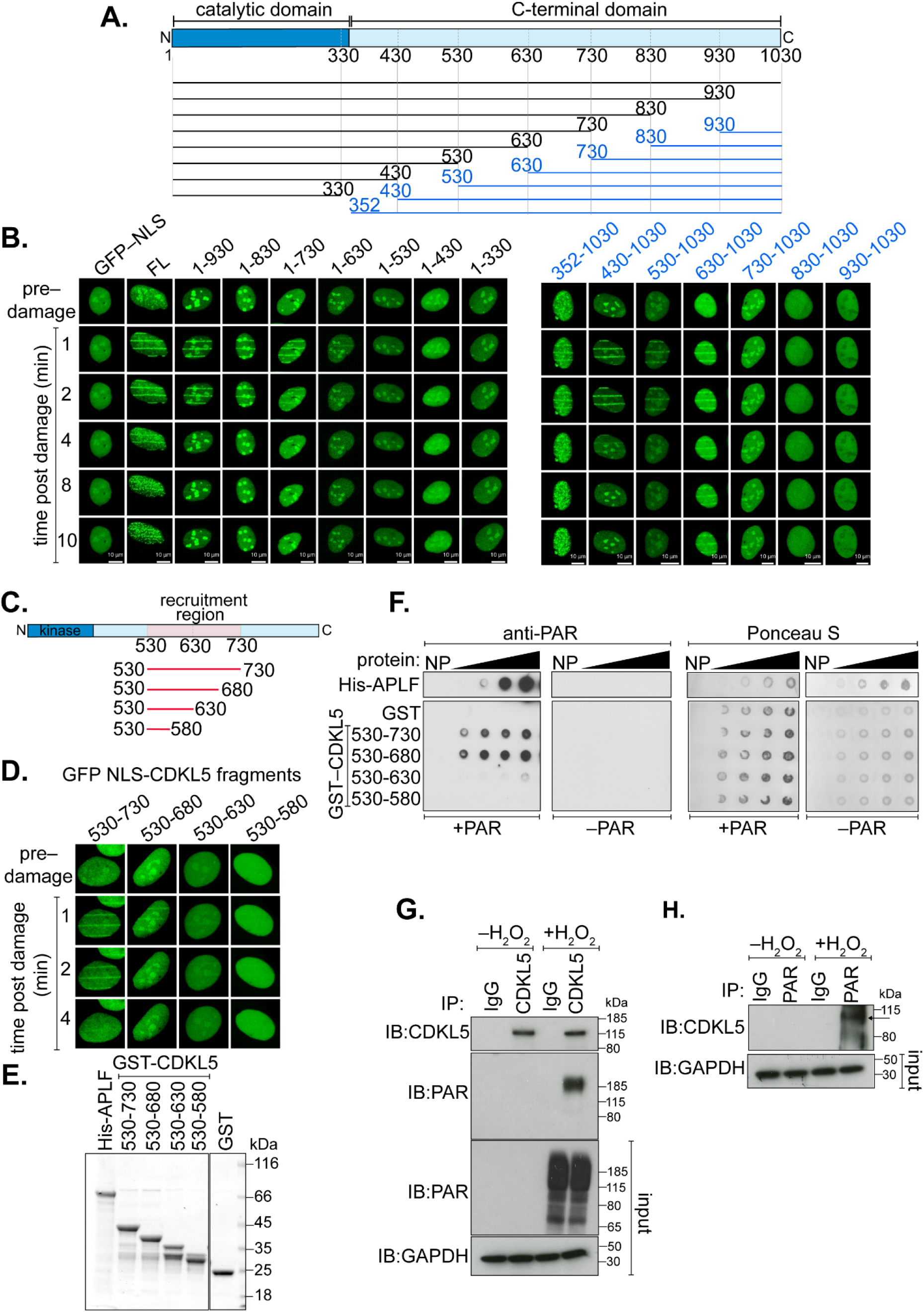
The CDKL5 recruitment domain binds PAR directly. (**A**) Schematic diagram of CDKL5 deletion mutants, deleting from the N–terminal (blue) or C–terminal (black) ends. All proteins were expressed with an N–terminal NLS and GFP tag. (**B**) BrdU–sensitized U–2–OS (Flp-In T-Rex) cells stably expressing GFP– NLS, the GFP–NLS–CDKL5 deletion mutants shown in A, or full length (FL) GFP–NLS–CDKL5 were subjected to line micro–irradiation (355 nm) and time lapse imaging. Three independent experiments were performed, and one representative experiment is shown. (**C**) Schematic for fragments corresponding to the PAR dependent recruitment region in CDKL5 as identified in B. (**D**) Same as in B except that the GFP–NLS tagged CDKL5 fragments indicated were examined. (**E**) Coomassie gel showing recombinant fragments of human CDKL5 fused to GST purified from bacterial lysates. GST and APLF were also purified as controls. (**F**) Recombinant fragments of CDKL5 fused to GST (1.2, 2.5, 5, 10 μg), or GST, were dot–blotted on nitrocellulose membrane and then incubated with synthetic PAR. PAR binding was detected by far western blotting. APLF was used as positive control. (**G**, **H**) U–2–OS (Flp-In T-Rex) cells stably expressing CDKL5 were either mock treated or treated with 500 μM H_2_O_2_ for 30 min. Extracts were subjected to immunoprecipitation with antibodies against CDKL5 (G) or PAR (H) (or non-specific IgG as control). Precipitates, and input lysates, were analyzed by western blotting using the indicated antibodies.

**Fig. S3.**
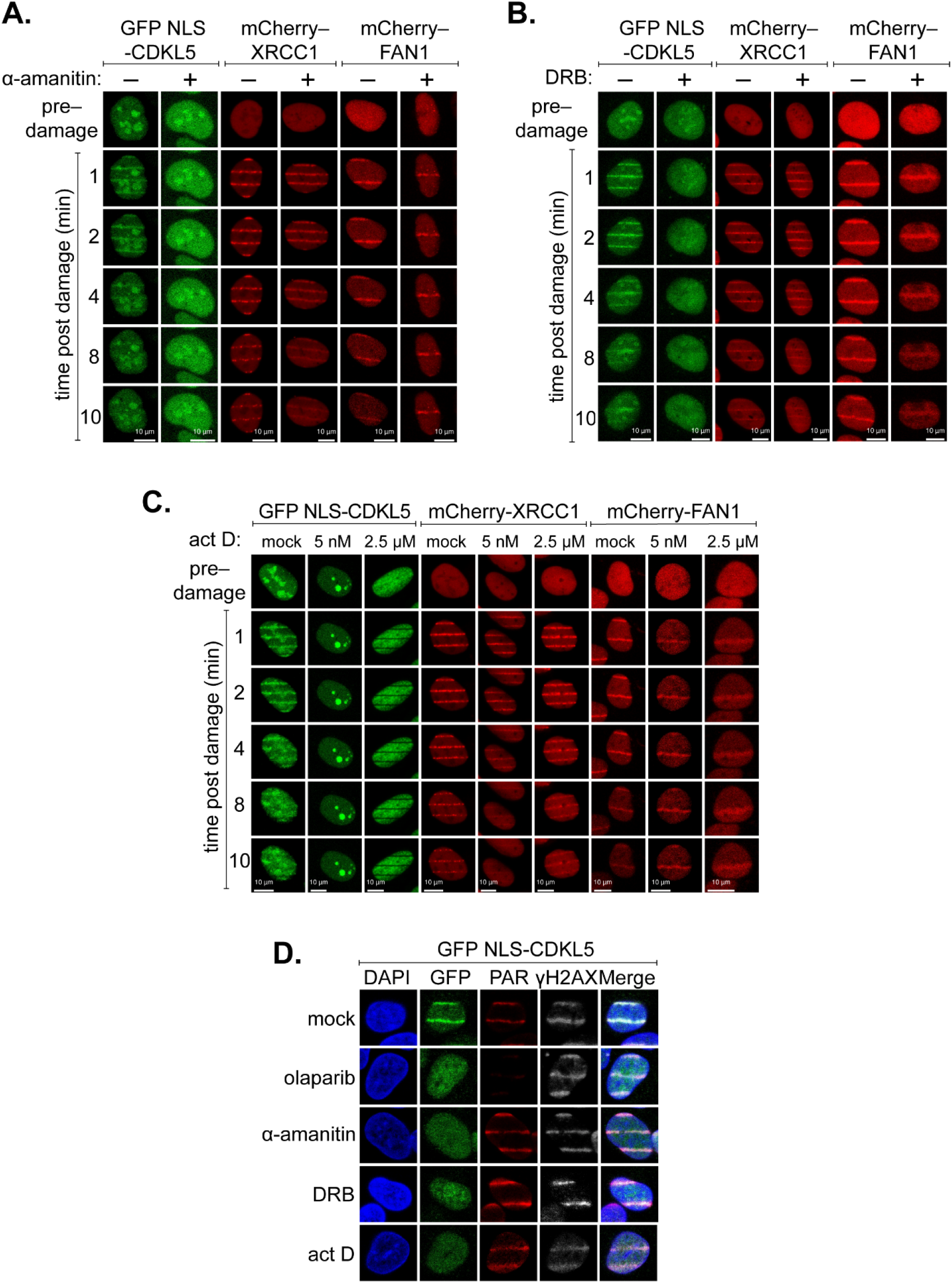
CDKL5 recruitment to DNA lesions requires ongoing transcription. (**A–C**) BrdU–sensitized U–2–OS (Flp-In T-Rex) cells stably expressing GFP–NLS–sCDKL5, mCherry–XRCC1 or mCherry–FAN1 were pre–incubated with α–amanitin (20 μg/ml, 8 h; A), DRB (100 μM, 2 hr; B) or actinomycin D (5 nM or 2.5 μM, 40 min; C) prior to line micro–irradiation (355 nm) and time lapse imaging. One of three independent experiments is shown. (**D**) Same as A–C. except that BrdU–sensitized cells stably expressing GFP–NLS–CDKL5 were also pre–incubated with olaparib (5 μM, 1h) as control. Cells were subjected to line micro–irradiation, fixed and then subjected to indirect immunofluorescence using antibodies against GFP, PAR and γH2AX.

**Fig. S4.**
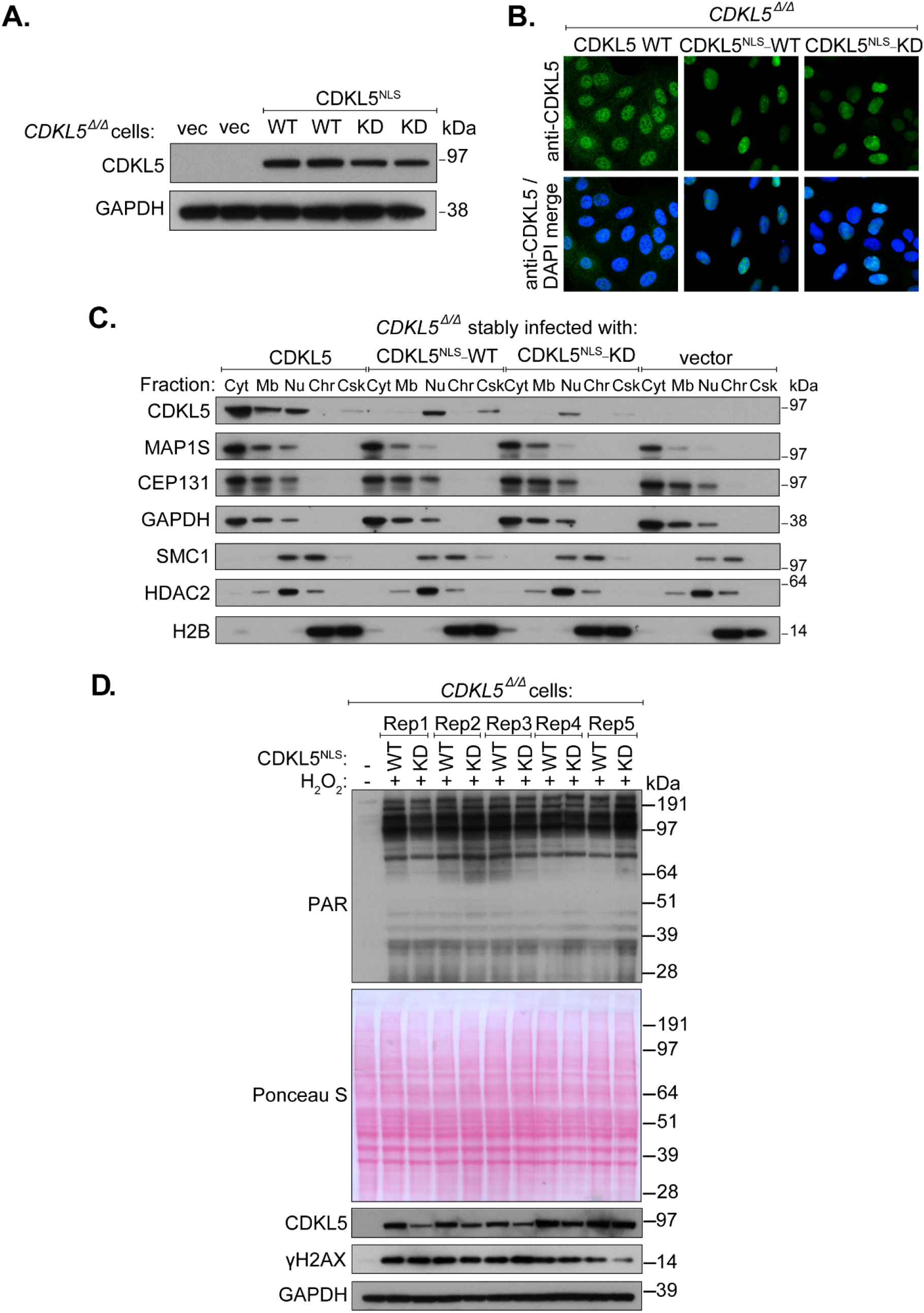
Restricting CDKL5 expression to the cell nucleus. (**A**) Extracts of CDKL5 disrupted U–2–OS (Flp-In T-Rex) cells (*CDKL5*^Δ/Δ^) stably expressing CDKL5^NLS^–WT or a K^42^R kinase-dead mutant (CDKL5^NLS^–KD) or empty vector were subjected to western blotting with the antibodies indicated. Two different dishes of cells are shown per condition. (**B**) *CDKL5^Δ/Δ^* cells stably expressing CDKL5, CDKL5^NLS^–WT or CDKL5^NLS^–KD were subjected to indirect immunofluorescence analysis with anti-CDKL5 antibodies. (**C**) Subcellular fractionation of lysates from *CDKL5*^΄/Δ^ cells stably expressing CDKL5, CDKL5^NLS^–WT or CDKL5^NLS^–KD or empty vector. Lysates were fractionated to isolate proteins found in the following subcellular compartments: cytoplasmic (Cyt), membrane (Mb), nuclear (Nuc), chromatin (Chr) or cytoskeleton (Csk). Fractionated samples were resolved by SDS-PAGE and probed with antibodies shown. (**D**) *CDKL5*^Δ/Δ^ cells stably expressing CDKL5^NLS^–WT or CDKL5^NLS^–KD (or empty vector) were treated with 500 μM H_2_O_2_ for 15 min. Samples were resolved by SDS-PAGE and probed with indicated antibodies or stained with Ponceau S to show equal loading. Rep=biological replicate.

**Fig. S5.**
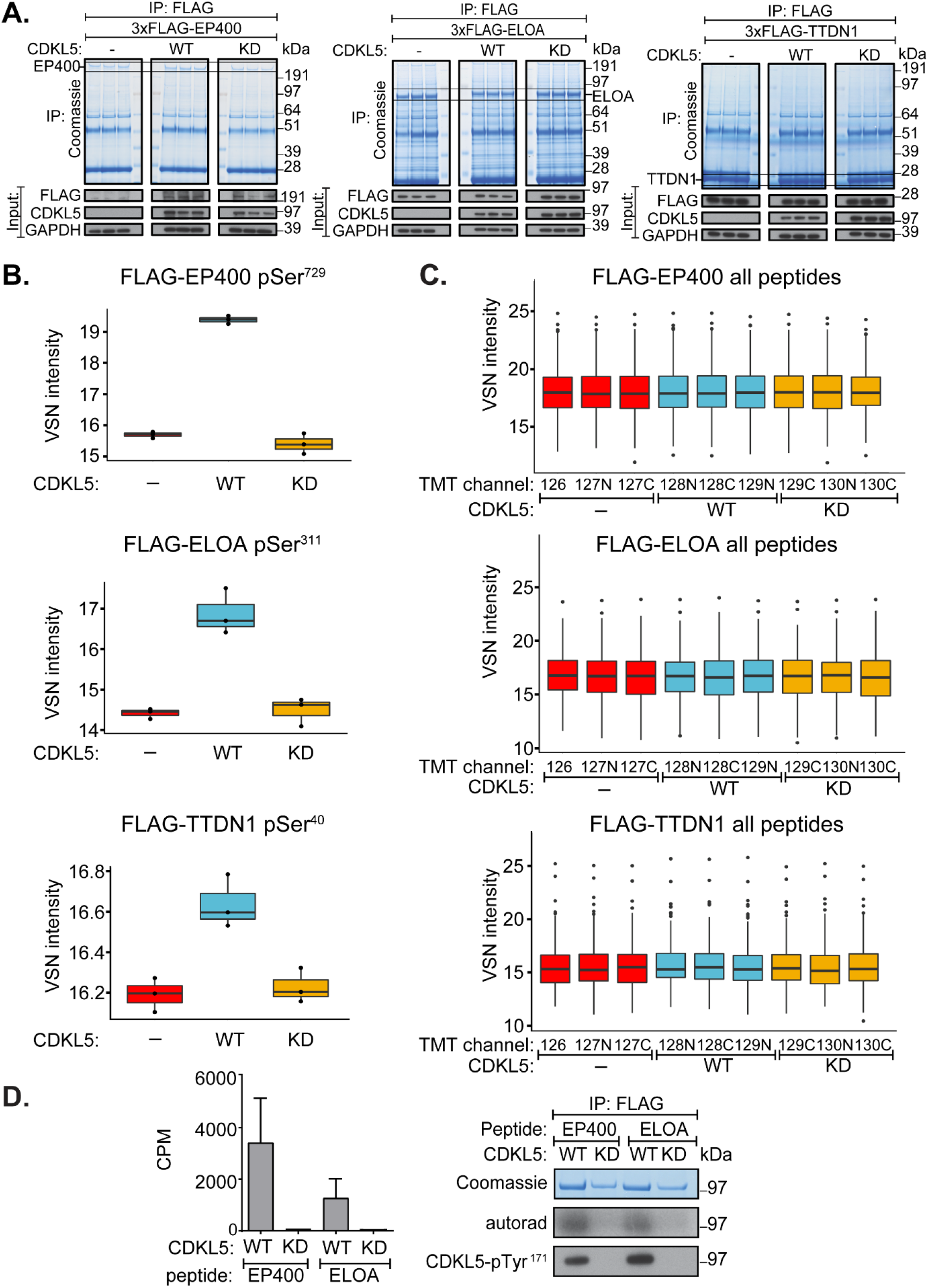
Validating phosphorylation of EP400, ELOA and TTDN1. (**A**) HEK293 cells were co-transfected with CDKL5^NLS^ (wild type “WT” or kinase-dead “KD” K^42^R mutant) and either FLAG-EP400 (left), FLAG-ELOA (middle) or FLAG-TTDN1 (right). 24 hr later cells were incubated with H_2_O_2_ (500 μM) for 15 min before being harvested and lysed. Protein extracts were subjected to immunoprecipitation with anti-FLAG-agarose beads. Precipitates were subjected to SDS-PAGE and blotting with antibodies shown (bottom panels) or staining with Coomassie Brilliant Blue (top panels). The bands corresponding to the FLAG-tagged proteins were excised from the gels in A. and processed for mass spectrometric detection of relevant phospho-peptides. Three independent co-transfection experiments were done for every condition. (**B**) Boxplots showing VSN–normalised intensity of phospho-peptides corresponding to EP400 pSer^729^, ELOA pSer^311^and TTDN1 pSer^40^ from the experiment in A. (**C**) Boxplots of the VSN-adjusted TMT reporter ion intensities for all peptides for each TMT label in the case of FLAG–EP400, FLAG–ELOA and FLAG–TTDN1 from the experiment in A. (**D**) *Left*: Anti-FLAG precipitates from HEK293 cells transiently expressing FLAG-tagged CDKL5 (wild type “WT” or a K^42^R kinase-dead “KD” mutant) were incubated with the synthetic peptides indicated, in the presence of [γ-^32^P]-labelled ATP-Mg^2+^ and peptide phosphorylation was measured by Cerenkov counting. Data are represented as mean ± SEM from three independent experiments. *Right*: Same but anti-FLAG precipitates were subjected to SDS-PAGE and autoradiography to detect CDKL5 autophosphorylation, or western blotting with CDKL5-pTyr^171^ antibody specific for the CDKL5-Tyr171 autophosphorylation site (*2*).

**Fig. S6.**
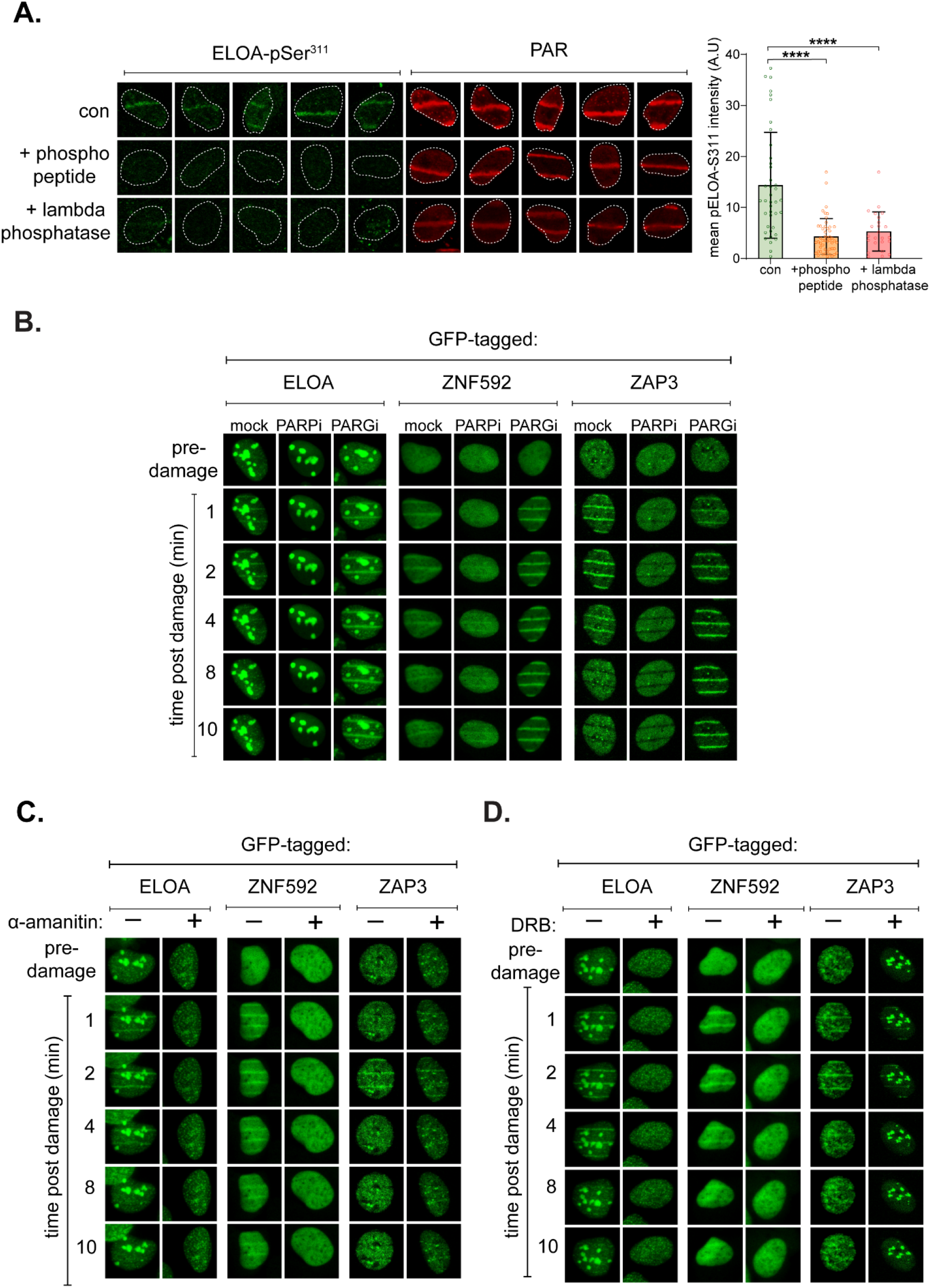
Recruitment of CDKL5 substrates to DNA damage sites. (**A**) BrdU–sensitized U–2–OS (Flp-In T-Rex) cells were subjected to nuclear line micro–irradiation (355 nm). Cells were fixed and then mock treated (con) or treated with lambda phosphatase prior to incubation with the primary antibodies indicated. Alternatively, ELOA-pSer^311^–phosphopeptide was included during incubation with the primary antibodies, before indirect immunofluorescence analysis. Quantification of ELOA-pSer^311^ signal at the laser tracks is shown. Data represent mean ± SD of two independent experiments. Statistical significance was assessed by one-way-ANOVA-test. Asterisks **** indicate *P*–values of <0.0001. (**B–D**) BrdU–sensitized U–2–OS (Flp-In T-Rex) cells stably expressing GFP–tagged forms of the proteins indicated were pre– incubated with olaparib (PARPi; 5 μM) or PD00017273 (PARGi; 0.3 μM, 1h) (B), α– amanitin (20 μg/ml, 8h) (C) or DRB (100 μM, 2 hr) (D) prior to line micro–irradiation (355 nm) and time lapse imaging. One of three independent experiments is shown.

**Fig. S7.**
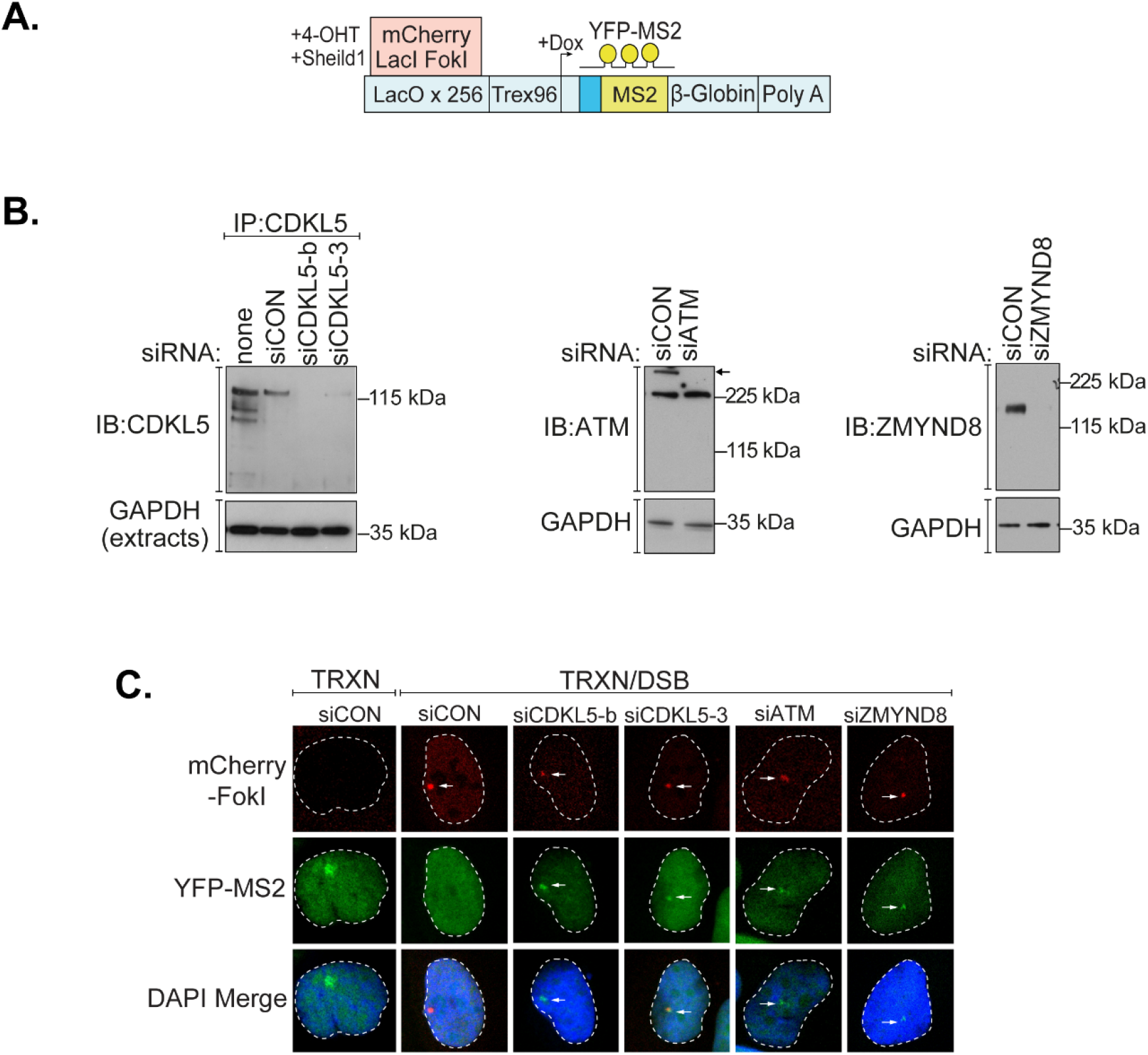
CDKL5 influences transcriptional activity at DNA breaks. (**A**) Cartoon of reporter construct (*3*) in which induction of the mCherry-tagged FokI endonuclease results in double-strand break (DSB) induction in a region upstream of a doxycycline-inducible reporter gene. Ongoing transcription of the reporter gene can be visualized by the presence of a YFP-MS2 fusion protein that binds stem-loop structures in the nascent transcript. (**B**) siRNA-mediated depletion of CDKL5, ATM and ZMYND8 in U–2–OS cells (263 IFII) harbouring the cassette shown in A. (**C**) U–2–OS cells (263 IFII) were transfected with the siRNAs indicated; siCON – non-targeting control. After addition of doxycycline to induce transcription, transcriptional silencing was monitored in cells before and after induction of the FokI endonuclease by quantification of YFP-positive cells. 150 cells were analysed per condition per experiment. Representative images of reporter cells are shown. Arrow indicate site of FokI mediated DSB (mCherry) and YFP-MS2 transcript.

**Movie S1.** Live–imaging of CDKL5 recruitment to sites of line–micro–irradiation. U–2– OS Flp–In T–REx cells stably expressing GFP–NLS–CDKL5 were preincubated with BrdU overnight. An hour before micro-irradiation along a line in the nucleus using a 355 nm laser attached to a Leica TCS SP8X confocal microscope, cells were mock treated or treated with PARP inhibitor (olaparib, 5μM, 1hr) or PARG inhibitor (PDD00017273, 0.3 μM, 1hr). Cells were live imaged for the time indicated.

**Movie S2. Live–imaging of CDKL5 recruitment to sites of spot–micro–irradiation.** U–2–OS Flp–In T–REx cells stably expressing GFP–NLS–CDKL5 were preincubated with BrdU overnight. Cells were mock treated or treated with PARP inhibitor (olaparib, 5 μM) or PARG inhibitor (PDD00017273, 0.3 μM) an hour before micro-irradiation in a spot-shaped sub-nuclear volume in the nucleus using a 405 nm laser attached to a Zeiss Axio Observer Z1 spinning disk confocal microscope. Cells were live imaged for the time indicated.

**Table S1. CDKL5 nuclear phospho-proteomics data.** List of the proteins with phosphopeptides that are more abundant in cells expressing CDKL5^NLS^–WT versus KD, with a fold change greater than 1.5. The mass spectrometry nuclear phosphoproteomics data have been deposited to the ProteomeXchange Consortium via the PRIDE (*4*) partner repository with the dataset identifier PXD022916.

**Table S2. Lists of the reagents, antibodies, plasmid constructs, siRNA sequences, peptide sequences and primer sequences used in this study.** Datasheets for each plasmid used in this study will be available on a dedicated page of our reagents website upon publication.

## Materials and Methods

### Reagents

All reagents including antibodies, cDNA clones, oligonucleotides and peptides used in the present study are enlisted in Table S2. After publication, all cDNA clones and antibodies generated in-house, and datasheets for each plasmid, can be requested via the MRC PPU Reagents and Services reagents website.

### ELOA phospho-Ser^311^ antibodies

ELOA-pSer^311^ antibodies were raised by MRC-PPU Reagents and Services at the University of Dundee in sheep and purified against the relevant antigen: (DA081; 3rd bleed; raised against the peptide KEENRRPPS*GDNARE conjugated to bovine serum albumin). Sheep were immunised with the peptide antigen followed by 4 further injections 28 days apart, with bleeds performed seven days after each injection.

### Cell lines and cell culture

All cells were kept at 37 °C under humidified conditions with 5% CO2. HEK293, HEK293T and U–2–OS Flp-In T-Rex, U–2–OS 263 IFII reporter cells were grown in GIBCO DMEM media (Life Technologies, Paisley, UK) supplemented with 100 U/ml penicillin, 100 μg/ml streptomycin, 1% (v/v) l-glutamate (GIBCO, Invitrogen), 1% (v/v) sodium pyruvate and 1% (v/v) non-essential amino acids, 10% (v/v) foetal bovine serum or 10% (v/v) TET System–approved FCS for U–2–OS reporter cell lines (631106, Takara Bio). U–2–OS-pEP15 cells (*5*) were maintained in 1 mg/ml glucose phenol-red-free DMEM (Lonza) supplemented with steroid-free FBS (Thermo Fisher Scientific), 1% antibiotic-antimycotic (Sigma-Aldrich), GlutaMAX™-I (Gibco) and 800 μg/ml G-418 (Sigma-Aldrich). U–2–OS (Flp-In T-Rex) cells were maintained in 10 μg/ml blasticidin. Hygromycin (100 μg/ml) or puromycin (2 μg/ml) were used to select for the integration of constructs in Flp-In recombination sites. All cell lines were regularly tested for mycoplasma contamination. U–2–OS Flp-In T-Rex *CDKL5*^Δ/Δ^ cells were described previously (*2*).

### Cell transfections

HEK293 cells were transfected using the calcium phosphate transfection protocol as previously described (*2*). Cells were seeded at a confluence of 20–30% in 15 cm plates and 24 hr later were co–transfected with a total of 10 μg of plasmid (5 μg + 5 μg in the case of plasmid co-transfection). Cells were incubated with the transfection mixture for 24 hr before being harvested and lysed.

For transient expression of GFP–tagged proteins in U–2–OS cells, cells were transfected with 1–2 μg of pcDNA5 FRT/TO plasmids using GeneJuice Transfection Reagent (Millipore) on to 1×10^5^ adhered U–2–OS or U–2–OS Flp-In T-Rex cells in 2 mL media in a 3.5 cm glass bottom dish (FD35–100, WPI). 8 hr following transfection, cells were incubated overnight with 0.5–1 μg/ml tetracycline hydrochloride to induce expression of the target protein.

For siRNA mediated knockdown of proteins, cells were transfected with a 100 nM suspension of relevant siRNA duplexes (Eurofins) or siRNA SMARTpools (Dharmacon) using Lipofectamine RNAi-MAX transfection reagent (13778150, Invitrogen, Paisley, UK) as per manufacturer’s guidelines. Cells were analyzed 60–72 hr following transfection. siRNA sources and sequences are outlined in **Table S2.**

### Generation of stable cell lines using the Flp-In T-REx systems

To generate U–2–OS (Flp-In T-Rex) cells stably expressing target proteins, cells were co-transfected with 9 μg of POG44 Flp-recombinase expression vector (Thermo Fisher) and 1 μg of pcDNA5 FRT/TO-target protein, using GeneJuice Transfection Reagent (Millipore). 48 hr following transfection, cells were selected in the presence of 100 μg/ml hygromycin and 10 μg/ml blasticidin in the medium. Around 10 to 12 days later, surviving colonies were pooled together and resulting cultures were analysed for the expression of target protein following induction with increasing amounts of tetracycline hydrochloride (T3383, Sigma-Aldrich). To generate cells stably expressing nucleus–restricted CDKL5, *CDKL5*^Δ/Δ^ cells (*2*) were infected with retroviruses expressing wild type CDKL5 with an exogenous nuclear localisation signal (CDKL5^NLS^–WT), a kinase dead (K^42^R) CDKL5^NLS^–KD or an empty vector. Thirty–six hours later, medium was replaced with fresh medium containing 3 μg/ml puromycin and cells were kept for another 48 hr in selection conditions and pooled together. A list of plasmid constructs is included in **Table S2**.

### Whole–cell extract preparation and western blotting

Cell pellets were lysed on ice for 30 min in ice–cold lysis buffer (50 mM Tris/HCl (pH 7.4) buffer containing 0.27 M sucrose, 150 mM NaCl, 1% (v/v) Triton X–100, 0.5% (v/v) Nonidet NP-40 and 0.1% (v/v) 2–mercaptoethanol) supplemented with a protease inhibitor cocktail (cOmplete™, EDTA–free Protease Inhibitor Cocktail), benzonase (Novagen, 50 U/ml), microcystin-LR (Cat. Number, 33893, Sigma) at a final concentration of 10 ng/ml and phosphatase inhibitor cocktail-2 (P5726, Merck) at 1% (v/v). The lysate was cleared by centrifugation at 17,000 *g* for 15 min and supernatant was collected for protein measurement by Bradford assay and storage at –80 °C. For western blotting, whole cell extract (40 μg) was mixed with LDS–PAGE sample buffer (Thermo Fisher) containing 5% (v/v) 2–mercaptoethanol before boiling at 95 °C. Samples were resolved by 4–12% Bis–Tris SDS–PAGE gradient gels (NuPAGE, Thermo Fisher) followed by transfer onto a Hybond–C Extra Nitrocellulose membrane (GE1060000, GE Healthcare) for 105 min at 100 V. The membrane was blocked in 5% (w/v) non–fat dry milk in TBS-Tween–20 (0.2% v/v) for 1 hr and probed with diluted primary antibodies. The membrane was washed three times in TBS–Tween–20 (0.1%(v/v)), incubated with secondary antibodies diluted in blocking buffer for 1 hr, washed three times in TBS–Tween–20 (0.1% (v/v)) prior to developing the membrane using SuperSignalTM West Pico PLUS Chemiluminescent Substrate (Thermo) and capturing the signal on an X-ray film. See **Table S2** for antibody and dilution information.

### In vitro phosphorylation of peptides using CDKL5 immunoprecipitates

HEK293 cells were transiently transfected with either wild-type or kinase dead (K42R) CDKL5-FLAG expressing constructs, and 48 hr later plates were washed in cold PBS and lysed in ice-cold buffer (50 mM HEPES (pH 7.5), 1% (v/v) Triton X-100, 0.27 M sucrose, 300 mM NaCl) freshly supplemented with protease inhibitor cocktail (cOmplete™, EDTA-free) 10 mM iodoacetamide, 10 ng/ml microcystin-LR, 2% (v/v) phosphatase inhibitor cocktail 2 (Sigma-Aldrich) and 500 U/ml universal nuclease. Lysates were then cleared by centrifugation for 10 min at 20,000 *g* at 4 °C and protein concentration was measured. Extracts (~2.0 mg) were then incubated with 10 μl (settled) anti-FLAG agarose M2 affinity beads (Sigma-Aldrich) for 2 hr at 4 °C. Beads were washed five times in lysis buffer containing 1 M NaCl and then twice in kinase buffer (50 mM Tris 7.5, 10 mM MgCl2, 0.1 mM EGTA). Beads were resuspended in 15 μl kinase buffer containing 0.15 mM peptide substrate and 0.1% (v/v) 2-mercaptoethanol. Reactions were initiated with the addition of 5 μl [γ-^32^P]-ATP (0.1 mM), incubated for 30 min at 30 °C with constant shaking and stopped by adding 10 μl of 0.5 M EDTA. Samples were centrifuged at 20,000 *g* and supernatants (30 μl) were spotted onto P81-phosphocellulose paper. Papers were then washed 5 times in 75 mM orthophosphoric acid, once in acetone and dried. ^32^P incorporation in each sample was measured by Cerenkov counting using a Perkin Elmer TriCarb Scintillator counter. Beads were resuspended in LDS sample buffer, boiled and subjected to SDS–PAGE followed by Coomassie staining.

### Laser micro–irradiation

#### “Line” micro–irradiation

Around 1×10^5^ cells expressing (stably or transiently) fluorescently tagged protein were seeded in 3.5 cm glass bottom dishes (FD35–100 for 24 hr in media containing 10 μM bromodeoxyuridine (BrdU–Sigma) and 0.5–1 μg/ml tetracycline hydrochloride (Sigma). Shortly prior to irradiation, cells were washed with PBS and the medium was replaced with warm, low absorption medium (31053, Thermo). Cells were placed in an incubator chamber at 37 °C with 5% CO2 supplementation mounted on a Leica TCS SP8X microscope system (Leica Microsystems). Laser micro–irradiation was performed using a protocol adapted from (*6*). Briefly, a striation pattern was generated by scanning bi–directionally at either 16×16 or 32×32 pixel resolution using a 355 nm laser (Coherent), resulting in a pattern of 16 or 32 horizontal lines across the imaging field. The laser dose was adjusted by altering the laser scanning speed and the number of scanning iterations per line. Typically, irradiation was performed by scanning at 5 Hz with 3 iterations per line. The power at the objective (approximately 1.5 mW) was measured using a power meter (Thorlabs). Using the above settings, we typically irradiated at approximately 1.4–2.8 J/m^2^. Laser micro–irradiation experiments were performed using a Leica HC PL APO CS2 63x/1.20 water objective, using a pre–defined imaging template utilizing the ‘Live Data Mode’ module within the Leica LASX software. After software–mediated autofocus, a pre–irradiation image was recorded, followed by 355 nm laser micro– irradiation. Time–lapse imaging was performed following the field of view every 30s for 5-10 min. Pre– and post– irradiation images were taken at 1024×1024 pixel resolution, scanning at 467Hz, taking eight 1 μm optical sections per image with 2x averaging. Pre– and post–irradiation images were stitched using an ImageJ macro and used for visualisation and analysis.

#### “Spot” micro-irradiation

Cells were prepared for imaging as described above. Cells were placed in an environmental chamber at 37 °C with 5% CO2 attached to an Axio Observer Z1 spinning disk confocal microscope (Zeiss). Micro–irradiation was performed using a single-point scanning device (UGA-42 firefly, Rapp OptoElectronic). Single–point regions of interest (ROI) were defined for each cell and irradiated with 100% 405 nm laser power for 600 iterations after removal of the ND filter. The estimated power delivered per ROI on average was approximately 27 J/m2. ROI x–y co–ordinates were recorded and used for subsequent image analysis. A pre–defined imaging template was used within the Zen Blue acquisition software. A pre–irradiation image was recorded, followed by 405 nm irradiation. A time–lapse was subsequently performed every 5 s for 10 min. Hardware autofocus (Definite Focus, Zeiss) was used to ensure focus was maintained throughout the time–lapse and was applied every 70 frames. To avoid image acquisition during laser micro–irradiation, a 3 s delay was applied from the start of micro–irradiation and the beginning of the time–lapse. Images were acquired using a C13440 camera (Hamamatsu), using a C Plan APO 64x/1.40 oil objective, acquiring 4x 0.5 μm optical sections per image with 4×4 binning.

#### Image analysis

Recruitment to sites of spot micro-irradiation was quantified using CellTool by modifying analysis protocol adapted from (*7*). Briefly, pre– and post–irradiation images were first stitched using an ImageJ macro. Maximum intensity projections of the stitched images were then taken. Individual cells were manually cropped from the original image and a 5×5 Gaussian blur filter applied to minimize the impact of noise on subsequent image processing. Micro–irradiated spots were then tracked using the spot–detector /track module within CellTool. Recruitment was calculated as the difference between the average intensity in the recruitment region and of a nearby region, multiplied by the total area of recruitment. For negative results, where the protein of interest was not recruited, ROI co–ordinates were imported to CellTool and the maximum recruitment within the static ROI was determined, as described above.

#### Drug treatment

PARP inhibitors Olaparib (S1060, Selleck chem) and Talazoparib (S7048, Selleck chem) and PARG inhibitor PDD00017273 (5952, Tocris bioscience) were used at a final concentration of 5 μM, 50 nM and 0.3 μM respectively, and were added to the cells 1 hr prior to and during micro–irradiation. Transcription inhibitors were employed as follows: α–amanitin (20 μg/ml) for 8 hr; DRB (100 μM) for 2 hr; actinomycin D (5 nM and 2.5 μM) for 40 min prior to, and for the entire duration of micro–irradiation. For RNAse treatment, cells were first washed with warm PBS and permeabilised with Tween–20 (1%(v/v)) in PBS for 5 min followed by treatment with 1 mg/ml RNase A (Thermo) for 10 min at room temperature (RT). Following the respective treatments, cells were micro–irradiated and imaged immediately.

### Immunofluorescence

Cells grown on coverslips were washed twice with cold PBS and fixed with 4% paraformaldehyde (sc–281692, Santa Cruz) in PBS for 15 min at RT. After fixation, cells were washed twice with PBS, and permeabilized with 0.5% (v/v) Triton–X–100 (in PBS) for 15 min at RT, washed twice with PBS and blocked at least 1 hr in antibody dilution buffer (1x PBS containing 5% Normal Donkey Serum, 0.1% (v/v) fish skin gelatin, 0.1% (v/v) Triton–X–100, 0.05% (v/v) Tween–20). Incubation with the relevant primary antibody (overnight at 4 ° C) was followed by three washes (5 min in PBS+0.05% (v/v) Tween–20) and incubation with appropriate fluorescently–labelled secondary antibody (60 min, RT). Coverslips were washed three times (5 min in PBS+0.05% (v/v) Tween– 20), stained with DAPI (Sigma) (1 μg/ml in PBS, 5 min) and mounted using ProLong Gold anti–fade mounting agent (P36934, Thermo).

To measure chromatin retention of CDKL5 after oxidative DNA damage, U–2–OS Flp-In T-Rex cells expressing GFP–NLS or GFP NLS–CDKL5 were grown on coverslips in media containing 1 μg/ml tetracycline. After 18 hr cells were pre–incubated with PDD00017273 (0.3 μM; “PARGi”) either in the absence or presence of PARP inhibitor olaparib (15 μM) for 60 min before exposing the cells to hydrogen peroxide (H1009, Sigma) (500 μM) for 30 min. Cells were then washed twice with cold PBS (containing 0.3 μM PARGi) and pre–extracted in cold 0.2% (v/v) Triton X–100 (in PBS containing 0.3 μM PARGi) for 4 min at room temperature prior to fixation as above. Imaging of fixed samples was carried out on a Leica TCS SP8 MP microscope using oil immersion objective (HPA CL APO CS2 63x/1.40 Oil). Quantification of detergent–insoluble anti– GFP signal (excluding nucleolar GFP signal) from >150 cells per sample per repeat were done using Fiji ImageJ based macro. Non–nucleolar anti–GFP fluorescence signal was quantified in the region co–localizing with DAPI but excluding the nucleolar region defined by fibrillarin co–labelling. Mean nuclear GFP fluorescence was plotted relative to that in untreated WT cells. Data were plotted and analysed by GraphPad Prism v9.0.0 using one-way ANOVA followed by Bonferroni multiple comparison test.

To examine phosphorylation of Elongin A at sites of DNA damage, 1×10^5^ U–2–OS Flp-In T-Rex cells (wild type, CDKL5 disrupted (*CDKL5*^Δ/Δ^) or cells pre-depleted with indicated siRNA for 48 hr) were seeded in 8 well chamber slides (Ibidi), 24 hr prior to the experiment, in media containing 10 μM Bromodeoxyuridine (Sigma). 0.3 μM PARG inhibitor (PDD00017273) was added to cells 30 min before the irradiation. Nuclei were irradiated as described previously. The cells were pre-extracted with cold 0.2% (v/v) Triton–X–100 (in PBS) for 2 min at RT and washed twice with cold PBS. Cells were fixed with 4% paraformaldehyde in PBS for 10 min at RT. After fixation, cells were washed twice with PBS, and permeabilized with 0.2% (v/v) Triton–X–100 (in PBS) for 5 min at RT, washed twice with PBS and blocked for 45 min in antibody dilution buffer (1x PBS containing 5% (v/v) Normal Donkey Serum, 0.1% (v/v) fish skin gelatin, 0.1% (v/v) Triton–X–100, 0.05% (v/v) Tween–20). Incubation with 0.32 μg/ml ELOA-pSer^311^ antibody (pre-incubated with 4.8 μg/ml corresponding non-phospho peptide for 12 hr at 4 °C) was done overnight at 4 °C, followed by three washes (5 min in PBS+0.05% (v/v) Tween-20) and incubation with appropriate fluorescently–labelled secondary antibody (60 min, RT). Cells were washed three times (5 min in PBS+0.05% (v/v) Tween-20), stained with DAPI (1 μg/ml in PBS, 5 min) and mounted using ProLong Gold antifade mounting agent. The buffers used in each step were supplemented with 1% (v/v) phosphatase inhibitor cocktail-2 and PhosSTOP (Roche: 1 tablet per 10 ml). Imaging of fixed samples was carried out on a Leica TCS SP8 MP microscope using oil immersion objective (HP CL APO CS2 63x/1.40 Oil). Treatment with olaparib and DRB was done prior to irradiation as explained before. To confirm the phospho-specificity of the ELOA-pSer^311^ antibody, fixed and permeabilised cells were (i) mock treated or treated with 100 U lambda phosphatase (NEB) overnight at 30 °C prior to primary antibody incubation (ii) incubated with the 0.32 μg/ml ELOA-pSer^311^ antibody that was pre-incubated with 6.4 μg/ml phosphopeptide for 12 h at 4 °C.

Quantification of ELOA-pSer^311^ to DNA damage sites was performed using a Cell Profiler image analysis pipeline. After segmentation and cropping of individual nuclei, micro-irradiation tracts delineated by PAR were segmented. Within each nucleus, the background nuclear intensity outside the segmented tracts was subtracted from the mean intensity from all detected irradiation tracts. Nuclei in which the background intensity was higher than the intensity within the micro-irradiation site were excluded. Data were plotted and analysed by GraphPad Prism v9.0.0 using one-way ANOVA followed by Dunnett’s multiple comparison test. The image analysis scripts are available on request.

### Recombinant protein expression and purification

Escherichia coli BL21 codon plus (DE3) cells transformed with expression plasmids encoding GST–tagged CDKL5 fragments (530–730, 530–680, 530–630, 530–580), or GST alone or His_6_–APLF were grown in Luria Broth (LB) medium containing 100 μg/ml ampicillin to *A*_600_ 0.5, followed by 0.5 mM isopropyl β-D-thiogalactopyranoside (IPTG) induction in early log phase for 16 hr at 20 °C. Cells were harvested by centrifugation at 3,500 g, and pellets were resuspended in lysis buffer (50 mM Tris–HCl pH 7.5, 150 mM NaCl, 5 mM DTT, 5 mM EDTA, 1 mM PMSF and 0.2 mg /ml lysozyme, 25 units Universal nuclease (Pierce™ Universal Nuclease for Cell Lysis), left on ice for 30 min followed by brief sonication on ice (5 cycles of 30 s on, 30 s off at 30% amplitude). The homogenate was centrifuged at 20,000 *g* for 30 min at 4 °C and the clarified cell lysates were applied to respective affinity resin columns. (i) The clarified cell lysates from cells overexpressing GST fusion proteins were applied to glutathione sepharose resin pre–equilibrated with equilibration buffer (50 mM Tris–HCl pH 7.5, 150 mM NaCl, 5 mM DTT and 5 mM EDTA, 0.1% (v/v) Triton X–100). The column was washed five times with equilibration buffer and twice with equilibration buffer without detergent. The GST– fusion proteins were eluted with 20 mM reduced glutathione in 50 mM Tris–HCl pH 7.5, 150 mM NaCl, 10% (v/v) glycerol, 1 mM DTT and 5mM EDTA. (ii) The clarified cell lysates obtained from cells overexpressing His_6_–APLF were applied to Ni–NTA resin pre–equilibrated with equilibration buffer (50 mM Tris–HCl pH 7.5, 150 mM NaCl and 5mM DTT, 0.1% (v/v) Triton X–100, 10 mM Imidazole). The column was washed five times with equilibration buffer and twice with equilibration buffer without detergent. The His_6_–APLF was eluted using 300 mM imidazole in 50 mM Tris–HCl pH 7.5, 150 mM NaCl, 10% (v/v) glycerol, 1mM DTT. Eluted proteins were dialysed overnight at 4 °C in sucrose buffer (50 mM Tris/HCl pH 7.5, 150 mM NaCl, 270 mM sucrose, 0.1 mM EGTA, 0.03% (v/v) Brij–35, 0.1% (v/v) β–mercaptoethanol). The proteins were concentrated, snap–frozen and stored at −80 °C for further use.

### In vitro poly–ADP ribose (PAR) binding assay

Serial dilutions (10, 5, 2.5, and 1.25 μg) of GST, GST–CDKL5 fragments and His_6_– APLF were dot–blotted onto an activated nitrocellulose membrane under low vacuum conditions. The membranes were dried and stained with Ponceau S to check loading. The membrane was washed and blocked with 5% (w/v) skimmed milk powder in PAR binding buffer (20 mM Tris/HCl pH 7.5, 50 mM NaCl) for 1 hr prior to incubation with 50 nM synthetic PAR (Trevigen) (in blocking buffer, 45 min, RT). The membrane was washed twice with PAR binding buffer followed by incubation with primary antibodies (rabbit anti–PAR polyclonal (Trevigen), 1:5000 in blocking buffer, 4 °C, overnight), and secondary antibodies (goat anti–rabbit HRP–conjugated; Thermo), 1:5000 in milk, 1 hr, RT. PAR binding buffer was used to rinse the membrane three times after each antibody incubation. The membrane was developed using SuperSignalTM West Pico PLUS Chemiluminescent Substrate (Thermo) and the resulting signal was captured on an X–ray film.

### Immunoprecipitation: CDKL5 binding to PAR in cells

U–2–OS Flp-In T-Rex cells stably expressing CDKL5 were mock treated or treated with H_2_O_2_ (500 μM; 30 min) in the presence of PDD00017273 (0.3 μM). CDKL5 was immunoprecipitated from 2 mg extract using anti–CDKL5 antibody (S957D); sheep IgG (31243, Thermo) was used as control. Cells were lysed in 50 mM Tris/HCl (pH 7.4) buffer containing 0.27 M sucrose, 150 mM NaCl, 1% (v/v) Triton–X–100, 0.5% (v/v) Nonidet NP–40 and 0.1% (v/v) 2–mercaptoethanol supplemented with a protease inhibitor cocktail (cOmplete™, EDTA-free Protease Inhibitor Cocktail), benzonase (Novagen, 50 U/ml), 10 ng/ml microcystin-LR (33893, Sigma), phosphatase inhibitor cocktail–2 (P5726, Merck) at 1% (v/v), 0.3 μM PARGi PDD00017273 (5952, Tocris bioscience) and 5 μM PARPi Olaparib. Extracts were then incubated for 30 min at 4 °C and clarified by centrifugation at 17,000 *g* in a refrigerated centrifuge. Clarified extracts were pre-cleared using DynaBeads Protein G (10003D, Life Technologies) conjugated with sheep IgG isotype control using manufacturer’s protocol, for 45 min at 4 °C. Pre–cleared extracts were used to immunoprecipitate CDKL5 using sheep polyclonal CDKL5 antibodies or sheep IgG isotype control with DynaBeads Protein G. Approximately 2 μg of anti–CDKL5/sheep IgG isotype control was linked to beads to perform pull down from 2 mg of pre–cleared extracts for 2 hr at 4 °C. Alternatively, pre-cleared extracts (2 mg) were incubated for 4 hr at 4 °C with 2 μg of pan–ADP–ribose binding reagent (MABE1016, Merck) or normal rabbit IgG (2729S, Cell Signalling) conjugated to DynaBeads Protein G. Beads were washed three times with lysis buffer and twice in cold PBS before boiling at 95 °C in LDS–PAGE sample buffer (Thermo Fisher) containing 5% (v/v) 2–mercaptoethanol. Samples were resolved in 4–12% Bis–Tris SDS–PAGE gradient gels (NuPAGE, Thermo Fisher). Input lysates or immunocomplexes were analysed by western blotting using sheep polyclonal anti–CDKL5, pan–ADP–ribose binding reagent and anti–GAPDH (14C10, Cell Signalling) antibodies. Antibodies were diluted in 5% (w/v) skimmed non-fat dry milk in TBS-Tween–20 (0.2% v/v). Membranes were incubated overnight at 4 °C or 2 hr at RT with the relevant antibodies, then washed. Membranes were then incubated with recombinant protein G–HRP (1: 2500) (ab7460) for 1 hr at RT. The membrane was developed using SuperSignalTM West Pico PLUS Chemiluminescent Substrate (Thermo) and the resulting signal was captured on an X–ray film.

### Phosphoproteomic identification of nuclear substrates of CDKL5

Twenty 15 cm plates of CDKL5 disrupted U–2–OS cells (*CDKL5*^Δ/Δ^) expressing CDKL5^NLS^–WT or CDKL5^NLS^ –K^42^R were grown to around 70% confluence, treated with H_2_O_2_ (500 μM for 15 min), washed twice with PBS and were harvested in 4 mL of ice–cold solution containing 20% (v/v) TCA, 80% (v/v) acetone and 0.2% (w/v) DTT, transferred into 5 mL Eppendorf tubes and stored at –20 °C overnight. Samples were then centrifuged twice at 20,000 *g*, –10 °C for 20 min and supernatants were then discarded. Pellets were resuspended with 2 mL ice–cold 80% (v/v) acetone and then centrifuged again at 20,000 *g*, –10 °C for 30 min. After removing the supernatants completely, pellets were left to air–dry for 10 min.

TCA/acetone–precipitated pellets were resuspended in 500 μL 8 M urea, 50 mM AmBiC, 1% (v/v) phosphatase inhibitor cocktail–2, 0.1% (v/v) microcystin, pH 8.0 and Benzonase at a concentration of 0.2% (v/v) and incubated 15 min at room–temperature, and finally lysed using a Bioruptor sonicator. Lysates were centrifuged at 20,000 *g* for 30 min at RT and stored at –80 °C for further mass spectrometry analysis. Five independent biological replicates were carried out, on different days.

Protein concentrations were determined using a BCA assay kit and measured the absorbance at 560 nm. A total of 5 mg protein from each sample were reduced with 5 mM DTT at 45 °C for 30 min, alkylated with 10 mM iodoacetamide at room temperature in the dark for 20 min), quenched by addition of 5 mM DTT, digested with Lys–C (1:200 (w/w), Lys–C:protein) for 4 h at 30 °C, then diluted with 50 mM ammonium bicarbonate to 1.5 M final urea concentration, and finally followed by trypsin digestion (1:50 (w/w), trypsin: protein) at room temperature overnight. 1% TFA (v/v) was added to stop the digestion. The acidified digests were centrifuged at 10,000 *g* for 10 min. The collected supernatants were then desalted on 200 mg Sep–PAK tC18 cartridges, and the eluents were dried by speed vacuum centrifugation (Thermo). Desalted peptides were resuspended in 1 ml of 2 M lactic acid, 50% (v/v) acetonitrile (ACN) and centrifuged at 15,000 *g* for 20 min. Supernatants were transferred to an Eppendorf tube containing 18 mg of titanium dioxide (TiO2) beads (GL sciences, Japan) and vortex–mixed for 1 hr at room temperature. The TiO2 beads were washed two times (10 min per wash) with 2 M lactic acid, 50% (v/v) ACN followed by three washes with 0.1% (v/v) TFA, 50% (v/v) ACN. Phospho–peptides were eluted twice with 150 μl of 10% (v/v) ammonia solution (NH_4_OH) and were finally eluted with 150 μl of 50% (v/v) ACN, 5% (v/v) ammonia solution (NH_4_OH). The combined eluent was dried with vacuum centrifugation and then cleaned up using in–house–made C18 StageTips (3M Empore™). 1% of each TiO2– enriched sample was analyzed by mass spectrometry prior to following processes.

TMT–10plex labeling was performed according to the manufacturer’s protocol using the TMT Labeling Kit. Briefly, The TiO_2_–enriched sample was resuspended into 100 μl of 100 mM TEAB. A total of 0.4 mg of each TMT tag were used for labeling each sample. After 1 hr incubation, 2 μl of each labeled sample was diluted with 18 μl of 0.1% formic acid and were then checked for TMT labeling efficiency. After checking the labeling efficiency, each TMT–labeled sample was quenched by incubation with 8 μl of 5% (w/v) hydroxylamine for 30 min at RT. The quenched samples were then mixed and fractionated with high pH reverse phase C18 chromatography using the Ultimate 3000 high–pressure liquid chromatography system (Dionex) at a flow rate of 569 μl/min using two buffers: buffer A (10 mM ammonium formate, pH 10) and buffer B (80% ACN, 10 mM ammonium formate, pH 10). Briefly, the desalted TMT labeled samples were resuspended in 200 μL of buffer A (10 mM ammonium formate, pH10) and fractionated on a C18 reverse phase column (4.6 × 250 mm, 3.5 μm, Waters) with a gradient as follow: 3% buffer B to 12.5 % buffer B in 5 min, 12.5% to 40% buffer B in 35 min, 40% B to 60% B in 15 min, 60% B to 100% B in 5 min, 100% for 5 min, ramping to 3% B in 5 min and then 3% for 10 min. A total of 60 fractions were collected and then concatenated into 20 fractions, which were further desalted over C18 StageTips and speed vacuum dried prior to LC–MS/MS analysis.

#### LC–MS/MS mass spectrometry

LC–MS/MS analysis was performed with an Orbitrap Fusion Lumos (Thermo), with a Thermo Dionex Ultimate 3000RSLC Nano liquid chromatography instrument. Peptide concentration from each fraction was quantified by Nanodrop, samples were dissolved in 0.1% formic acid, and 1 μg of each fraction was loaded on C18 trap column with 3% (v/v) ACN, 0.1% (v/v) TFA at 5 μl/min flow rate. Peptides were separated over an EASY–Spray column (C18, 2 μm, 75 μm x 50 cm) with an integrated nano electrospray emitter (flow rate 300 nl/min). Peptide separation was done over 180 min with a segmented gradient applying following buffer system: Buffer A: 0.1% (v/v) formic acid, Buffer B: 80% (v/v) acetonitrile, 0.08% (v/v) formic acid. The first 7 fractions started from 6%–35% buffer B in 120 min (Note: the following 7 fractions started from 8% and the last 6 fractions started from 10%), 35%– 45% buffer B in 30 min, 45%–95% buffer B for 5 min, followed by 5 min 95% in buffer B. Eluted peptides were analyzed on an Orbitrap Fusion Lumos (Thermo Fisher Scientific, San Jose, CA) mass spectrometer. Spray voltage was set to 2.2 kV, RF lens level was set at 30%, and ion transfer tube temperature was set to 275 °C. The Orbitrap Fusion Lumos was operated in positive ion data–dependent mode with HCD fragmentation and orbitrap detector for all precursor fragments for reporter ion quantitation. The mass spectrometer was operated in data–dependent Top speed mode with 3 s per cycle. The full scan was performed in the range of 350–1500 *m/z* at nominal resolution of 120,000 at 200 *m/z* and AGC set to 4×10^5^ with maximal injection time 50 ms. The MS2 scan was set with an isolation width of 1.2 *m/z* with no offset, followed by selection of precursors above an intensity threshold of 5×10^4^ for Higher–energy Collisional Dissociation (HCD)–MS2 fragmentation with 38% normalized collision energy. Dynamic exclusion was set to 60 s. Monoisotopic precursor selection was set to *peptide*, maximum injection time was set to 120 ms. Charge states between 2 to 7 were included for MS2 fragmentation and analysis of fragment ions in the orbitrap using 50,000 resolving power with auto normal range scan starting from *m/z* 100 and AGC target of 5×10^4^.

#### Phosphoproteomics data analysis

Mass spectrometry raw data was searched against the Uniprot database (*homo sapiens*, including protein isoform sequences, 42,326 entries, downloaded 05/04/2018 from www.uniprot.org) using MaxQuant (version 1.6.3.4) (*8*). Variable modifications were set to: oxidation of methionine, phosphorylation of serine, threonine and tyrosine, deamidation of asparagine, carbamylation of the peptide N-terminus and acetylation of the protein N–terminus. Fixed modification was set as carbamidomethylation of cysteine. False discovery rate threshold for peptide identification was set to 5%. Quantitative results data was analysed using an in–house R (Version 4.0.0) (*9*) analysis pipeline (Script Files S1-S4). In brief, the intensities of peptides with more than one observation within a single sample fraction were averaged. Peptides quantified in several fractions were averaged in each respective fraction independently to avoid reporter ion quantification bias caused by differences in precursor co–isolation populations between different sample fractions. Data was normalized and calibrated using variance stabilizing normalization (VSN) (*10, 11*). Statistical testing was carried out using linear models for microarrays (*limma*) (*12*). under application of robust hyperparameter estimation (*13*). Peptides were declared significant based on visual assessment of a modified volcano plot (termed here “sprinkler” plot). Peptide metadata was extracted from the following databases: PhosphoSitePlus(R) (downloaded from www.phosphosite.org, version: 031120 (*14*), Uniprot (gene ontology (GO) data, downloaded from www.uniprot.org on 05/04/2020) and STRING (version 11.0, downloaded from https://string-db.org/). GO terms and protein–protein interaction networks were analysed using R (Script Files S1-S4). Following R packages were used: ggplot2 (*15*), reshape2 (*16*), vsn (*10*), *limma* (*12*), seqinR (*17*), plyr (*18*), stringr (*19*), ggrepel (*20*), ggpointdensity (*21*), wesanderson (*22*), extrafont (*23*), scales (*24*), matrixStats (*25*), GO.db,(*26*), STRINGdb (*27*), igraph (*28*) and ggnetwork (*29*). Session information is listed in Text File S1. The mass spectrometry nuclear phospho-proteomics data have been deposited to the ProteomeXchange Consortium via the PRIDE (*4*) partner repository with the dataset identifier PXD022916.

Data analysis Script Files (S1–S4), Session information text file S1, and the relevant database links can be downloaded from the Zenodo link: https://doi.org/10.5281/zenodo.4311475

### Extracted ion chromatogram (XIC) analysis

HEK293 cells were transfected using the calcium phosphate transfection protocol as previously described (*2*). Cells were seeded at a confluence of 20–30% in 15 cm plates and 24 hr later were co–transfected with 5 μg DNA for each plasmid. Cells were kept with the transfection mixture for 24 hr and then either mock treated or incubated with H_2_O_2_ at a final concentration of 500 μM for 15 min. Plates were then washed twice with phosphate saline buffer and lysed in a 50 mM Tris/HCl (pH 7.4) based buffer containing 0.27 M sucrose, 150 mM NaCl, 1% (v/v) Triton X–100, 0.5% (v/v) Nonidet NP–40, and 0.1% (v/v) 2–mercaptoethanol. Lysis buffer was freshly supplemented with a protease inhibitor cocktail (cOmplete™, EDTA–free Protease Inhibitor Cocktail), benzonase at 50 U/ml, microcystin–LR at 10 ng/ml final concentration, phosphatase inhibitor cocktail–2 (Merck) at 1% (v/v), olaparib (10 μM) and PDD00017273 (2 μM). Lysates were incubated for 30 min at 4 °C and clarified by centrifugation at 20,000 *g* at 4 °C.

For extracted ion chromatography (XIC) analysis, approximately 25 μl (settled volume) of FLAG–M2 agarose (Sigma–Aldrich; F1804) beads were mixed with the following amounts of lysate for 2–3 hr at 4°C: 10 mg of crude lysate for EP400 samples, 6 mg for Elongin A and 2.5 mg for TTDN1. Precipitates were then extensively washed with lysis buffer and finally once in cold PBS. Samples were then denatured in 25 μl LDS–PAGE sample buffer (Thermo Fisher) supplemented with 5% (v/v) 2– mercaptoethanol and then incubated at 95 °C for 5 min. All the immunoprecipitations were done in triplicates using lysates from independent replicate transfections. Samples were resolved in 4–12% Bis–Tris SDS–PAGE gradient gels (Nupage, Thermo Fisher) and relevant bands were excised and further processed for mass spectrometry as detailed below. Protein bands excised from the gel were destained, and proteins digested with Trypsin/LysC as described in (*2*). Peptides were labelled with TMT10plex (Thermo Fisher) according to manufacturer’s protocol, omitting the TMT131 label. Deviating from the protocol described in (*2*), labelling reaction was stopped using a 5% (w/v) hydroxylamine (Sigma) solution. After labelling, the peptides were freeze dried and stored at –80 °C until required.

Peptides were resuspended in 2% (v/v) acetonitrile (Merck), 0.1% (v/v) formic acid (Merck), incubated 15 min in an ultrasonic bath (VWR) and afterwards transferred into glass autosampler vials (Waters). Peptides were separated and analysed using the instrumental setup as described for the phospho–proteomics dataset. Elution of peptides was achieved by a segmented linear gradient over 120 min: Initial 3 min of isocratic 3% B, followed by 3% B to 7% B in 2 min, to 25% B in 60 min, to 45% B in 30 min, to 95% B in 5 min and isocratic state at 95% B for 5 min. This was followed by a linear gradient from 95% B to 5% B within 0.5 min and column re–equilibration for 14.5 min at 5% B. Flow rate was set to 300 nl/min. MS precursor ion scan was conducted within the Orbitrap at a resolution of 120,000 at 200 *m/z*. The top 15 precursors within a mass range of 350–1500 *m/z* were isolated in the quadrupole (0.7 Da isolation window, AGC target: 4×10^5^, max. injection time 50 ms) for subsequent fragmentation using HCD (38% normalized collision energy, AGC target: 5×10^4^, max. injection time 120 ms) and analysed in the Orbitrap with a resolution of 50,000 at 200 *m/z*. Analysed peptides were dynamically excluded after their first measurement from re–analysis for a duration of 60 s. Data was recorded in profile mode. In case of the file “Ivan_EP400–TMT.raw” an inclusion list of 628.3314 *m/z* (TMT labelled phospho-peptide SSPVNRPSpSATNK) was set. Orbitrap run metadata was extracted using the MARMoSET R package as described on their GitHub page (https://github.molgen.mpg.de/loosolab/MARMoSET, accessed 28/11/2020) (*30*).

Mass spectrometry raw data was searched using MaxQuant (Version 1.6.3.4). Variable and fixed modifications with the exclusion of carbamylation of the peptide N-terminus, FASTA, and FDR thresholds were set as described above for the phospho–proteomics dataset. Data was analysed using in–house written R scripts (Script Files S5-S8; Supplementary Information) which were modified from (*2*). In brief, all TMT reporter intensities of the identified peptides of the respective protein were normalized using VSN and intensities were statistically tested using a t–test with subsequent Bonferroni correction of the significance threshold of α = 0.05. Peptides with a p–value < 0.0125 (4 tests: EP400, TTDN1) or < 0.00833 (6 tests: ELOA) were considered significant. Session information is listed in Text File S2. The mass spectrometry extracted ion chomatography data have been deposited to the ProteomeXchange Consortium via the PRIDE (*4*) partner repository with the dataset identifier PXD022975. Data analysis Script Files and Session information text files, with links to the relevant databases can be downloaded from the Zenodo link: https://doi.org/10.5281/zenodo.4311494.

### U–2–OS FokI Transcription Reporter Assay

#### MS2 foci

U–2–OS 263 IFII transcription reporter cells (*3*) transfected with relevant siRNA were seeded on to 8–well chamber slides and treated with 1 μM Sheild1 (632189, Clontech Laboratories UK Ltd) and 1 μM 4–hydroxytamoxifen (4–OHT) (H7904–5MG, Sigma) for 3 hr to induce mCherry–FokI expression and 1 mg/ml doxycycline hyclate (D9891–G, Sigma–Aldrich) for an additional 3 hr to induce reporter gene transcription. Cells were fixed with 4% paraformaldehyde in PBS (sc–281692, Santa Cruz), washed three times with PBS, permeabilised with 0.2% (v/v) Triton–X/PBS for 3 mins at RT, washed and stained with DAPI (1 μg/ml in PBS, 5 min, RT) and mounted using ProLong Gold antifade mounting agent (P36934, Thermo). Imaging of fixed samples was carried out on a Leica TCS SP8 MP microscope using oil immersion objective (HAP CL APO CS2 63x/1.40 Oil). The number of transcription positive cells were scored manually from a total of 150–200 cells per variable in each independent repeat. Data were plotted and analysed by GraphPad Prism v9.0.0 using one-way ANOVA followed by Bonferroni multiple comparison test.

#### RNA Isolation, reverse transcription, and quantitative real-time PCR

RNA was extracted from 1.2×10^6^ cells using E.Z.N.A.® Total RNA Kit I (R6831–01) following manufacturer’s protocol. cDNA was synthesized from 1 μg RNA using iScript cDNA synthesis Kit (170–8891). qPCR was performed using a CFX384 real–time PCR system (Bio–Rad), relevant primers with 2% (around 20 ng) of the cDNA, and TB Green™ Premix Ex Taq™ II (Tli RNase H Plus) (RR820L, Takara) with two repeats for each PCR. The ΔΔC_t_ method was used for evaluation. GAPDH gene was used as a housekeeping gene for normalisation. Data were analysed in Excel software (Microsoft) and plotted in GraphPad Prism v9.0.0 software. Statistical significance was determined by two-way ANOVA followed by Dunnett’s multiple comparison test. Primers used are listed in Table S2.

### Gene silencing in response to I-PpoI-mediated DSB induction

2×10^5^ cells/ml/well U–2–OS-pEP15 cells (DOX-inducible ER-I-PpoI expressing stable cell line) were seeded into 6-well plates for siRNA transfection. Cells were transfected with 40 nM siSCR (Dharmacon) or 40 nM siCDKL5 (Dharmacon) using Interferin transfection reagent (Polyplus). Next day, 16 hr prior to the first 4–OHT treatment, 1 μg/ml DOX (Sigma-Aldrich) was applied to induce the expression of I-PpoI endonuclease. The following day, 1 μM 4–OHT (Sigma-Aldrich) treatment was used at different time-points (2, 4 and 8 hr) to facilitate the nuclear translocation of I-PpoI. 56 hr after siRNA transfection cells were collected and destined for RNA and gDNA isolation. For RNA isolation, ReliaPrep RNA Tissue Miniprep System (Promega), while for gDNA isolation, ReliaPrep gDNA Tissue Miniprep System (Promega) is used according to the manufacturer’s instructions. cDNA reverse transcription was performed with Applied Biosystems TaqMan Reverse Transcription Reagents (Thermo Fisher Scientific) according to the manufacturer’s instructions. qPCR experiments were performed on RotorGene Q qPCR machine. For data evaluation, ΔΔC_t_ calculation was used. Data were plotted and analysed by GraphPad Prism v9.0.0 using two-way ANOVA followed by Tukey’s multiple comparison tests.

## Notes

https://doi.org/10.5281/zenodo.4311475

https://doi.org/10.5281/zenodo.4311494

http://www.ebi.ac.uk/pride/archive/projects/PXD022975

http://www.ebi.ac.uk/pride/archive/projects/PXD022916

### Competing Interest Statement

The authors have declared no competing interest.

### Summary of Updates

The main article, figures, supplementary figures and supplementary information is now combined in a single pdf file.

